# Assessing the impact of renewable energy installations on biodiversity and identifying sustainable trade-offs

**DOI:** 10.64898/2026.02.24.707751

**Authors:** Marie-Ange Dahito, David Yang Shu, Gabriel Wiest, Stefano Moret, Tobias Wechsler, Loïc Pellissier

**Affiliations:** Swiss Federal Institute for Forest, Snow and Landscape Research (WSL), Zürcherstrasse 111, 8903, Birmensdorf, Switzerland; Department of Environmental Systems Science, ETH Zürich, Universitätstrasse 16, 8092, Zurich, Switzerland; Department of Mechanical and Process Engineering, ETH Zürich, Tannenstrasse 3, 8092, Zurich, Switzerland; Research Unit RECOVER, INRAE Aix-Marseille University, 3275 Route Cézanne, 13100, Aix-en-Provence, France; Department of Management, Economics and Industrial Engineering, Politecnico di Milano, Via Lambuschini 4/b, 20156, Milan, Italy

**Keywords:** biodiversity, renewable energy, hydropower, wind power, photovoltaic, energy systems planning, characterization factors

## Abstract

Renewable energy is crucial to achieve climate neutrality, but its rapid expansion can threaten biodiversity through habitat loss or fragmentation and ecological disruption. We present a spatially explicit assessment framework that quantifies biodiversity impacts from land use change associated with renewable energy infrastructure across a broad range of species groups, and identifies siting configurations that balance energy provision and conservation goals. Drawing on metrics from life cycle assessment, combined with species distribution models and siting strategies, we evaluate alternative deployment strategies. Using Switzerland as a case study, we compare three siting strategies (maximizing energy output, minimizing biodiversity impact, and a trade-off approach) for photovoltaic systems, run-of-river hydropower, and wind turbines. For solar and hydropower installations, prioritizing energy efficiency yields the highest cumulative biodiversity losses. However, these impacts can be substantially reduced with only a slight increase in land use by favouring biodiversity protection. For wind installations, strict avoidance of sensitive ecosystems may increase total impacts, as less efficient and therefore additional sites are required to achieve the same annual energy yield. Overall, our results show that trade-off-based siting strategies can effectively balance performance and biodiversity protection, highlighting that renewable energy can be provided without sacrificing sensitive ecosystems.

## 1 Introduction

Climate change and biodiversity loss are interconnected environmental challenges. Climate change disrupts species distributions, alters ecosystem dynamics, and erodes ecosystem services [IPBES et al., 2019, IPCC, 2023]. The resulting loss of biodiversity reduces ecosystem resilience and weakens forests, wetlands, and other natural carbon sinks, exacerbating climate change. Thus, both crises must be addressed at once.

While habitat destruction, pollution, and over-exploitation of resources have historically been the primary drivers of biodiversity erosion [Millennium Ecosystem Assessment, 2005], recent increase in the rate of climate change and the frequency of extreme events create additional pressures on biodiversity that will rival habitat loss in the second half of the century [Newbold, 2018]. Of nearly 170,000 species globally assessed by the International Union for Conservation of Nature (IUCN) Red List, over a quarter face extinction [IUCN, 2025] and more than 10,000 are under direct threat from climate change [IUCN, 2019]. Hence, greenhouse gas emissions must be cut rapidly as the main driver of climate change, including an energy transition based on renewable technologies such as hydro, solar, and wind power [IPCC, 2023, Bogdanov et al., 2019].

However, while renewable technologies can dramatically reduce the impacts of climate change, they may harm biodiversity if poorly sited [Katzner et al., 2013, Gasparatos et al., 2017]. Hydropower installations fragment river systems [Kuriqi et al., 2021, He et al., 2024] and alter flow regimes [Wechsler et al., 2023, Brunner and Naveau, 2023, Bätz et al., 2025] as well as the transport of sediments and organic matter [Arroita et al., 2015, Yu et al., 2019]; ground-mounted solar panels may compete with natural or agricultural areas [Hernandez et al., 2015, Zhang et al., 2024]; and wind energy installations affect flying species [Thaxter et al., 2017, Tolvanen et al., 2023]. Therefore, careful spatial planning based on ecological assessments is critical to ensuring a sustainable and biodiversity-friendly energy transition.

Measures have been taken to mitigate biodiversity loss, as is the case in the European Union with the adoption of the EU Birds and Habitats Directives. These two key laws led to the creation of Natura 2000, the largest network of protected areas worldwide [Evans, 2012]. Nonetheless, many ecologically valuable areas remain outside protected zones, due to incomplete national inventories, weak ecological connectivity, or inadequate management [Dubos et al., 2022, Castillo et al., 2020, Oberosler et al., 2020].

Siting decisions can be supported by spatial conservation planning tools such as *Marxan* [Ball and Possingham, 2000, Ball et al., 2009], *Zonation* [Moilanen et al., 2005], and *ConsNet* [Ciarleglio et al., 2009], which are designed to identify priority conservation areas. These tools rely on biodiversity data such as species distribution models (SDMs), habitat suitability indices, or range maps [Moilanen et al., 2009, Pressey et al., 2007]. However, even when socio-economic layers are included [Salak et al., 2024], these tools are not designed to directly provide implementable siting decisions for infrastructure. Their outputs represent relative spatial priorities, typically at the scale of planning units, and therefore do not incorporate fine-scale constraints required for infrastructure deployment. Besides, these methods do not quantify the specific biodiversity impacts of proposed infrastructure projects. Thus, comparisons between different siting options are not possible for supporting transparent decision making [McIntosh et al., 2018].

Life cycle assessment (LCA) is a methodology commonly used to quantify environmental impacts of human activities over the full life cycle. LCA methods specific to biodiversity impacts have been developed [Damiani et al., 2023], and can be used to support planning decisions. However, they commonly have low spatial resolution due to data limitations [De Baan et al., 2013, Verones et al., 2013, May et al., 2020], their underlying datasets tend to be biased toward vertebrates [Damiani et al., 2023], and they cannot be applied for local infrastructure siting decisions. Chaudhary et al. [2015] and Scherer et al. [2023] studied the biodiversity impacts of different land-use types and intensities on amphibians, birds, mammals, reptiles, and plants, with the latter study also considering land fragmentation. They both consider species vulnerability on a global level.

Specialized methodologies applied in LCA with higher spatial resolution are commonly limited to specific geographic regions (e.g., [Zelm et al., 2011, Geyer et al., 2010, de Baan et al., 2015, May et al., 2021]). Few studies quantified the biodiversity impacts of renewable energy installations on limited taxonomic groups. For example, May et al. [2021] investigated the effects of 39 existing onshore WT on the species richness of 13 bird species groups in Norway, and Dorber et al. [2020] evaluate the projected biodiversity impact per kWh for 1933 possible future reservoirs worldwide, considering five species groups, with fish being the only representative for aquatic biodiversity.

Neither of these studies explores siting strategies or proposes trade-off site selection frameworks. Furthermore, they consider only a limited number of taxonomic groups and rely mostly on low-resolution data, such as species–area relationship (SAR) models derived from global-scale studies, rather than on locally calibrated biodiversity information. To our knowledge, no existing work jointly evaluates land-use change and biodiversity impacts of renewable infrastructure expansion across a wide range of species groups, while providing a framework to identify siting configurations that balance electricity generation and conservation priorities.

Here, we address these limitations by developing a spatially explicit renewable energy siting framework that integrates LCA metrics with fine-resolution biodiversity modelling to evaluate biodiversity impacts associated with land use change across a wide range of taxonomic groups. We apply this framework to investigate the trade-offs between maximizing energy conversion and preserving biodiversity, considering three siting strategies. We demonstrate its capabilities by assessing the effects of the siting strategies for renewable installations on biodiversity impacts in Switzerland. In this case study, we consider the effects of projected expansions of photovoltaic (PV) panels, wind turbines (WT), and run-of-river (ROR) hydroelectric plants in 2050. We use high-resolution locally calibrated datasets, such as detailed species distribution and range data for 20 taxonomic groups, SARs tailored by species group and calculated at bioregion [FOEN, 2020] and country levels, region-specific extinction probabilities, and renewable energy potential maps. Thus, we account for both local and system-wide biodiversity impacts, as well as local renewable potentials and system-wide electricity demands. Our assessments consider installation-specific impacted species groups and extinction rates based on an extensive literature review (see Table 3), ensuring that siting decisions reflect technology-specific ecological pressures.

## 2 Methods

The proposed infrastructure siting framework integrates local biodiversity impact assessment through a spatially explicit data-driven approach. It requires spatial data on the state of biodiversity in the geography under investigation, spatial estimates of the expected energy conversion potential of renewable technologies, and the identification of exclusion areas where installation is undesirable or not possible. Based on these inputs, the framework quantifies local, regional, and global biodiversity impacts associated with land use change due to infrastructure deployment. Within the framework, alternative siting strategies can be developed and systematically assessed to identify options that balance energy provision potential with biodiversity conservation.

This framework is broadly applicable for assessing the biodiversity impacts of infrastructure siting in any geographic region, provided that suitable data are available.

### 2.1 Metrics for biodiversity impact due to land use change

To quantify the biodiversity impacts of land use change associated with infrastructure installation, we combine two metrics that are regional extinction probabilities (REPs)[Adde et al., 2025a] and characterization factors (CFs). The resulting metric measuring the overall biodiversity impact at the scale of the entire study region is hereafter referred to as the global CF.

The REP is based on the global extinction probability (GEP)[Kuipers et al., 2019], which assesses the risk of global extinction for a species group resulting from local extirpations and is computed for each region and species group based on species range sizes, threat levels, and spatial distributions.

Adde et al. [2025a] introduced the REP to preserve the high resolution of regional datasets and assess extirpation risks more accurately. The REP adapts the GEP to the spatial scale of the studied area.

For a species group 𝒢, the REP, whose value lies between 0 and 1, is computed per pixel as:

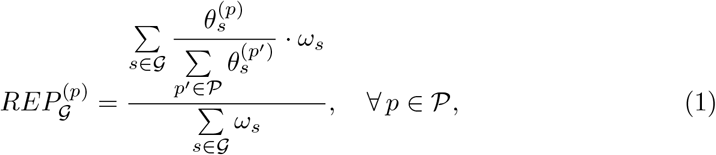

where 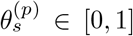 is the occurrence value of species *s* at pixel *p* based on species distributions (see Table 1), 𝒫 is the set of all pixels of the considered region, and *ω*_*s*_ ≥ 0 is the weighting factor representing the threat level of species *s* (see Table 2).

**Table 1:**
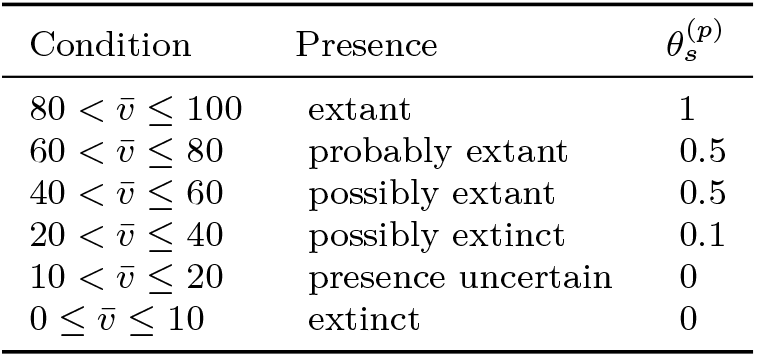
Weighting scheme for species occurrence 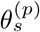.

| Condition | Presence | $\theta_s^{(p)}$ |
| --- | --- | --- |
| $80 < \bar{v} \leq 100$ | extant | 1 |
| $60 < \bar{v} \leq 80$ | probably extant | 0.5 |
| $40 < \bar{v} \leq 60$ | possibly extant | 0.5 |
| $20 < \bar{v} \leq 40$ | possibly extinct | 0.1 |
| $10 < \bar{v} \leq 20$ | presence uncertain | 0 |
| $0 \leq \bar{v} \leq 10$ | extinct | 0 |

**Table 2:**
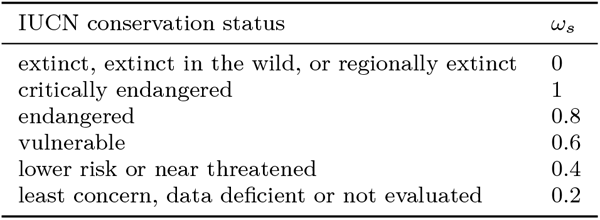
Linear weighting scheme associated to the IUCN threat level of each species *s*.

| IUCN conservation status | $\omega_s$ |
| --- | --- |
| extinct, extinct in the wild, or regionally extinct | 0 |
| critically endangered | 1 |
| endangered | 0.8 |
| vulnerable | 0.6 |
| lower risk or near threatened | 0.4 |
| least concern, data deficient or not evaluated | 0.2 |

**Table 3:**
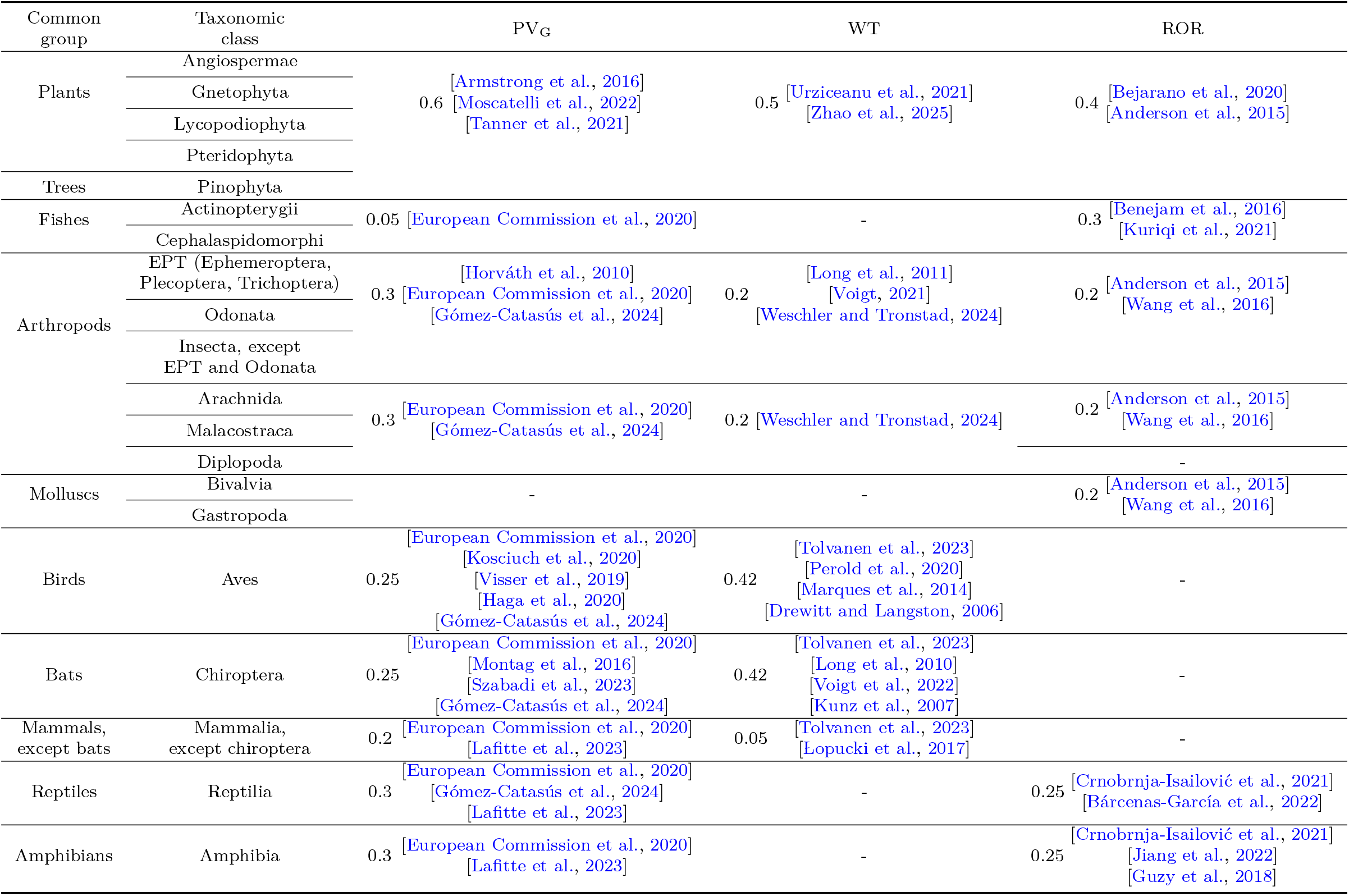
Species groups and associated extinction rates for ground-mounted photovoltaic, wind, and run-of-river power plants. Abbreviations: Ephemeroptera, Plecoptera, Trichoptera (EPT); ground-mounted photovoltaic (PV_G_); wind turbines (WT); run-of-river hydropower (ROR).

CFs are used in LCA to score environmental damage, allowing different pressures on ecosystems to be consistently assessed and compared. To translate biodiversity impacts associated with new infrastructure installations into measurable values, we adapt the methodology proposed by Chaudhary et al. [2015] and compute CFs using the pre-existing land use as the reference condition, instead of a natural reference state. This choice enables a specific focus on land-use change impacts (rather than land-use). We compute the CFs for spatial units defined as sets of pixels within the study region (e.g. a single pixel or all pixels within an installation area).

The loss of reference area due to land use change at spatial unit *u* ⊆𝒫 is:

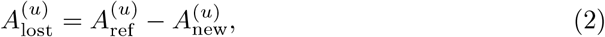

where 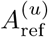 is the area under the reference land-use state and 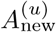 the area that remains unchanged after potential land-use modification. The regional loss in species richness due to new cumulative land use at spatial unit *u* is:

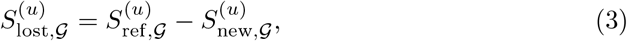

where 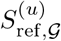 and 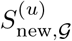 are the species richness of 𝒢 at the reference state and at the new state, respectively. 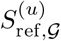 is derived from the species distribution data (see Section 2.3 and Supporting Information), while 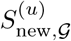 is obtained using the installation-specific extinction rate of 𝒢, which is represented by the local CF, denoted 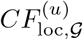, that is,

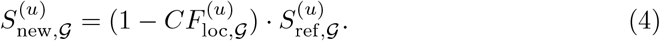

The derivative of the loss in species richness [Chaudhary et al., 2015] can be expressed as:

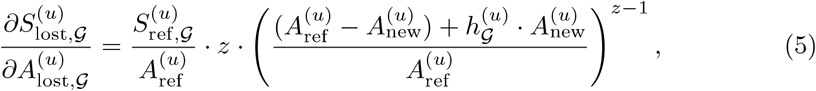

where 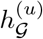 is the affinity of 𝒢 to the new land-use type, calculated as:

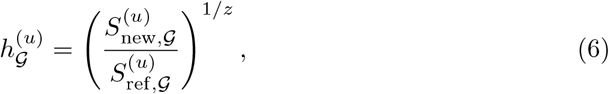

and *z* is the power parameter of the classic SAR of 𝒢.

The regional damages resulting from land occupation and transformation are expressed via a regional CF as the equivalent species loss throughout the operating lifetime of the installation (species-eq lost·years). The regional CFs for land occupation 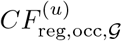 and land transformation 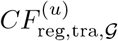 at spatial unit *u* ⊆ 𝒫 can be written as follows:

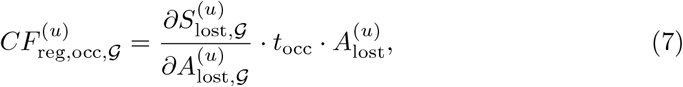

and

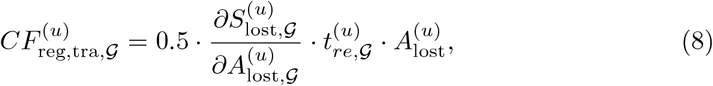

where *t*_occ_ is the duration for which the land is occupied by the installation, and 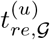 is the regeneration time needed to recover the biodiversity of the species group 𝒢 once all installations are removed and the land is not managed.

The overall regional impact 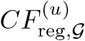 on group 𝒢 at spatial unit *u* is:

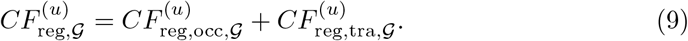

At the scale of the study region 𝒫, we characterize the global impact as the regional impact adjusted to reflect regional species vulnerabilities. The global impact on species group 𝒢 at spatial unit *u* can be written as follows:

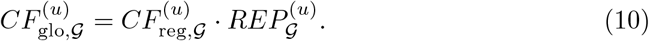

Notably, the probability of extinction at a spatial unit *u* is simply the sum of the extinction probabilities of its constituent pixels, that is,

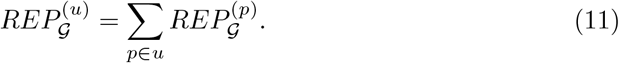

Multiplying the regional CF by the REP yields the global CF, expressed in species-eq lost·years, which accounts for the species ranges and vulnerabilities.

### 2.2 Approaches for biodiversity impact calculation

We use two complementary approaches to calculate biodiversity impacts (i.e., global CFs Equation (10)) from infrastructure deployment: a fine-scale pixel-level assessment and a global infrastructure-level assessment.

In the fine-scale approach, the equations described in Section 2.1 are applied at the pixel scale, that is, *u* ≡ *p* in Equations (2) to (11), with the parameter *z* specific to the bioregion in which each pixel is located (see SAR model calculation described in Section 2.3 and the Supporting Information). Therefore, CFs are calculated for every pixel. This method captures fine-scale spatial variation, allowing for a detailed understanding of how impacts differ across regions. It is particularly useful for identifying biodiversity hotspots and areas most affected by development (see the Supporting Information). However, to obtain a total global impact value, the impacts from all individual pixels of the study region must be aggregated. The choice of aggregation method does not alter the ranking of stressors, but affects the resulting values [Verones et al., 2015], which may influence the interpretation of the results and introduce additional complexity.

In contrast, to avoid aggregation effects, the global-level impact calculation treats the entire impacted zone as a single region of interest. In this case, the spatial unit *u* considered in Equations (2) to (11) corresponds to the set of all pixels impacted by new installations, *i*.*e*. the pixels containing the infrastructure together with their associated impact areas (see Supporting Information).

While it still incorporates local biodiversity information, such as the spatial distribution of selected pixels and combined SDM maps, this approach directly yields a single global CF which captures the overall effect of the infrastructure on the considered species group. Here, the *z* parameter is taken as the national scale estimate, and a single regeneration time is used, computed as the mean regeneration time across all impacted pixels. Finally, the REP is simply the sum of the REPs of the affected pixels.

The global-level approach does not provide insights into the spatial distribution of impacts, but avoids the need for aggregating pixel-scale values. As such, it is well suited for comparing overall biodiversity outcomes across different infrastructure siting scenarios. Therefore, we use the global-level impact calculation in Section 3 to generate biodiversity impact curves as a function of national energy output in our case study. Analyzing the impacts of renewable energy installations at a pixel scale enables to identify areas most vulnerable to infrastructure installation. The results for the pixel-level approach and additional Switzerland-specific analyses are provided in the Supporting Information.

### 2.3 Case study

We apply our framework to Switzerland, assessing the biodiversity impacts of siting four renewable energy technologies to meet installation targets required in the base scenario of the *Energy Perspectives 2050+* [Kemmler et al., 2021] for a net-zero energy system in Switzerland: rooftop and ground-mounted PV systems, WT, and small ROR hydropower plants.

#### Calculation of the extinction probabilities

For each species group 𝒢, we compute the REP values for 25m×25m-resolution pixels, considering only species ranges inside Switzerland.

We start from presence probabilities 0 ≤ *v* ≤ 100 (see Supporting Information) that are given for each pixel *p* and species *s*, obtained from species habitat suitability maps. A species is assumed to be possibly extant only when its presence probability in the pixel exceeds a presence threshold *τ*_*s*_ (see Supporting Information). To make the probabilities comparable across species, we follow the approach of Adde et al. [2025a] and scale the probabilities so that a value of 40 represents the lower bound for a species to be considered “possibly extant”, regardless of its original threshold:

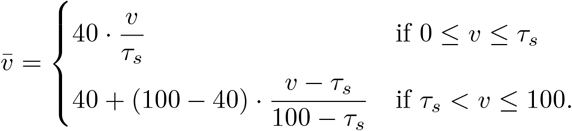

The species occurrence values 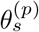 are obtained using the weighting scheme of Kuipers et al. [2019], which is based on Montesino Pouzols et al. [2014]. Table 1 shows the mapping of the scaled presence probabilities 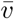 to the species occurrence values 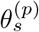.

Kuipers et al. [2019] compare three threat-level quantification schemes (linear, categorical, and logarithmic). Of the three schemes, we chose the linear categorization approach. Using Equation (1), we calculate the REP values.

#### Calculation of the regional characterization factors

In line with the REPs, we calculate CFs at a spatial resolution of 25m × 25m.

We use the most recent land cover data (see Supporting Information) to define the reference state prior to the installation of new energy infrastructure. The affinity of each species group 𝒢 (Equation (6)) is assumed to be equal to 0 if no species of 𝒢 is present at the reference state.

We assume the land occupation for the full lifetimes of the installations. We set the occupation times to 20 years for WT, 30 years for both rooftop and ground-mounted PV panels, and 80 years for ROR hydropower plants, based on average lifetimes reported by the life-cycle inventory database ecoinvent 3.11. These values are based on data for 2 MW onshore WT (at global scale), 3 kWp multi-Si PV flat-roof installation in Switzerland, 570 kWp multi-Si PV plant on open ground (at global scale), and ROR hydropower plants in Switzerland.

The regeneration times of species groups are based on [Curran et al., 2014] where average recovery times were computed for different families of species under passive and active restoration, distinguishing forest and non-forest biomes. We specifically use the predicted average recovery times relative to species richness for passive restoration in palearctic realm of latitude 45°. For the recovery of fish species, for which no data was available, we assume a recovery time of one year as these species can rapidly recolonize an aquatic area once an installation is removed. Details on the species groups considered are provided in the Supporting Information.

The parameter *z* (Equations (5) and (6)) was estimated by fitting SAR models to species distribution data using the R package `sars` [Matthews et al., 2019], with species data aggregated at the catchment level. Models were built for each bioregion and at the national scale. Details on the SAR models are provided in the Supporting Information.

#### Data collection

The case study assesses biodiversity impacts of renewable energy installations across Switzerland using national-scale geospatial datasets at a 25 m×25 m resolution, covering both terrestrial and aquatic ecosystems.

Biodiversity patterns are represented using fine-resolution SDM maps derived from SDMapCH [Adde et al., 2025c,b] integrated with a range mapping algorithm [Fopp, 2025] based on validated occurrence records (1980 – 2021) [Dépraz et al., 2025]. These data are used to generate binary presence–absence maps and species richness layers for the considered taxonomic groups (see Supporting Information).

Species groups potentially affected by land use change were identified for ground-mounted PV systems, WT, and small ROR hydropower based on the literature (see Table 3). Where available, published estimates of species richness reduction were used as local CFs; otherwise, assumptions were informed by expert judgment. Rooftop PV systems were assumed to have negligible biodiversity impacts, as they rely on existing infrastructure [Katzner et al., 2013].

Impacted areas were defined according to technology type and species mobility. For hydropower, impacted river sections were represented as polygons extended laterally to account for both aquatic and riparian effects. For PV and WT, minimum impacted areas correspond to ground coverage, with additional buffers applied for mobile species to reflect disturbance effects during operation. Buffer distances and their ecological justification are provided in the Supporting Information.

Production potentials of renewable energy technologies determine the number and spatial distribution of installations. We use data of the Swiss Federal Office of Energy [Klauser et al., 2022] to get the annual rooftop PV potential in Switzerland. Ground-mounted PV potential was derived from national solar irradiation datasets under current climate conditions [Sharma, 2023], accounting for panel efficiency, ground coverage, topography, and land-use exclusions. Based on [Dujardin and Lehning, 2022], wind energy potential was calculated on regional capacity factors and representative turbine models, while small hydropower potential relied on the national hydropower inventory [Hertach, 2012, SFOE, 2012] and the hydraulic potential model HYDROpot integral [Hirschi et al., 2013, Laub et al., 2022]. Detailed assumptions, exclusion criteria, and parameter values are documented in the Supporting Information. Across all technologies, areas of biodiversity importance were excluded in line with conservation guidelines. A complete list of protected-area datasets and exclusion buffers, as well as the treatment of border effects, is provided in the Supporting Information. The maps showing the production potential for ground-mounted PV, WT and ROR hydroelectric plants, along with the corresponding ERI computed at pixel level, are presented in Figure A1.

### 2.4 Siting strategies

We consider three siting strategies for selecting pixels where energy infrastructure can be installed. Pixels in excluded area or without energy generation potentials for the considered technology are not considered for siting. For ROR plants, the selected locations are computed as the centroids of the river section polygon geometries.

For a given energy infrastructure, we define the boolean variable *x* = [*x*_1_, *x*_2_,…, *x*_*n*_] ^⊤^, where *n* is the total number of suitable pixels. The elements of *x*, which are relative to each pixel, are only active (i.e., equal to 1) if the corresponding pixels are selected for the installation. The production potentials of pixels are given by *V* = [*V*_1_, *V*_2_,…, *V*_*n*_] ^⊤^.

Following Adde et al. [2025a], an extinction risk indicator (ERI) is derived by normalizing the logarithm of the REP:

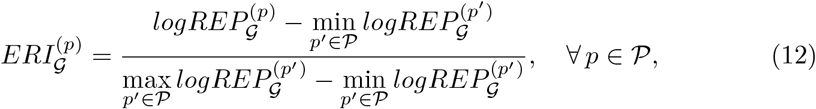

where 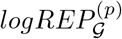 is the logarithmic transformation of 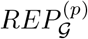 where zero values have been replaced before the transformation.

For all pixels, we determine the ERI *E* = [*ERI*_1_, *ERI*_2_,…, *ERI*_*n*_] for a combined extinction risk raster that accounts for all impacted groups. Therefore, 𝒢 in Equation (12) stands for the group consisting of all species impacted by the renewable energy considered. This normalized metric is particularly suited for trade-off strategies based on convex combinations to balance energy provision and extinction risk. The ERI is independent of the new infrastructure location, as it considers only biodiversity data in the reference state.

Finally, the demand *D* corresponds to the minimum production to be reached by the considered technology.

#### Maximizing production

The first strategy minimizes the surface transformed for the new installations, by choosing locations with the highest potential of production. We consider this approach as a proxy of the minimum cost strategy, since it reduces the number of required installations. The first siting approach is formalized as:

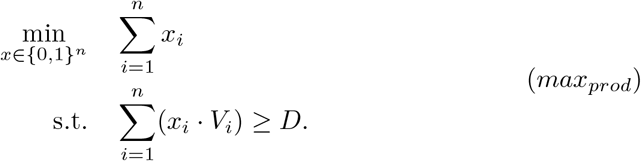

#### Minimizing extinction risks

The second strategy selects pixels with minimum risk of extinction for potentially impacted species groups, thus avoiding areas with high conservation values in terms of species richness of these groups. The ERI serves here as a biodiversity impact indicator, and the strategy can be formulated as:

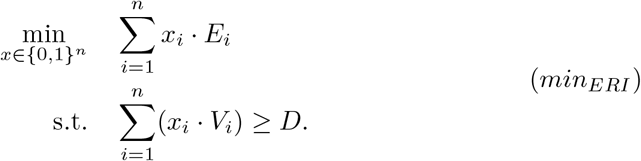

Here, we apply the ERI as the biodiversity impact indicator to reduce computation time, as the global CF can only be computed after all siting decisions have been made (Section 2), thus resulting in a computationally expensive combinatorial problem. To solve Equation (*min*_*ERI*_), the pixels are prioritized according to the lexicographic order (*E*_*i*_, −*V*_*i*_), that is, the strategy selects regions with the lowest ERI values, prioritizing higher-production areas when extinction risks are equal.

#### Identifying trade-offs

Finally, we consider a trade-off strategy that balances the surface area used and biodiversity impact. To that end, we introduce a weighting parameter *λ* ∈ [0, 1]:

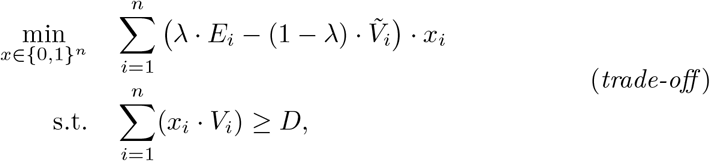

where 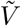 is the normalization of *V* in [0, 1].

We assess site selections, varying *λ* between 0 and 1. For each value, available pixels are prioritized based on the objective function defined in Equation (*trade-off*), resulting in unique siting configurations for each *λ*. As with the *min*_*ERI*_ approach, we apply a lexicographic sorting 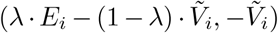: when multiple pixels share the same objective value from Equation (*trade-off*), those with higher production potential are prioritized.

Because the ERI serves only as a proxy for biodiversity damage in this selection strategy, we also compute the global CFs associated with each value of *λ* and its corresponding selected pixels. These are used to estimate a Pareto front describing joint trade-off between biodiversity impact and infrastructure size.

In the results, we apply the least-distance-to-ideal method to identify a representative trade-off siting. For this purpose, the global CFs and infrastructure sizes of the non-dominated solutions are normalized between 0 and 1. In this normalized two-dimensional space, the ideal siting corresponds to the minimum impact and minimum installation size, that is, the point (0, 0). We then select the Pareto solution with the smallest Euclidean distance to this ideal point, which we report as the trade-off siting configuration.

## 3 Results

The biodiversity impacts of rooftop PV, ground-mounted PV, WT and ROR in Switzerland are calculated on a global national level and for the local level on a pixel basis, as introduced in Section 2.2. In this section, we compare the results across various site-selection scenarios on a national level.

In the baseline scenario of *Energy Perspectives 2050+* [Kemmler et al., 2021] for a net-zero Swiss energy system, PV energy conversion in Switzerland increases by 29.34 TW h between 2025 and 2050, and the projected expansion of production from WT of 4 TW h. In our calculations of energy infrastructure deployment, we use these expansions as production goals for both rooftop and ground-mounted PV systems, as well as WT, accounting for local solar irradiation and wind availabilities. For small hydroelectric plants, an expansion of 0.77 TW h is assumed in the baseline scenario, mainly involving micro-hydropower plants, each with an installed capacity of less than 300 kW. To estimate the additional hydroelectric capacity required, we used a median number of full load hours of 4397 h per year, based on Swiss hydropower plant statistics [Dasen and Hertach, 2012]. Using this metric, an additional 182 MW of capacity is required to produce the target 0.77 TW h per year. A summary of the results presented in this section is provided in Table 4, covering all energy types and siting strategies.

**Table 4:**
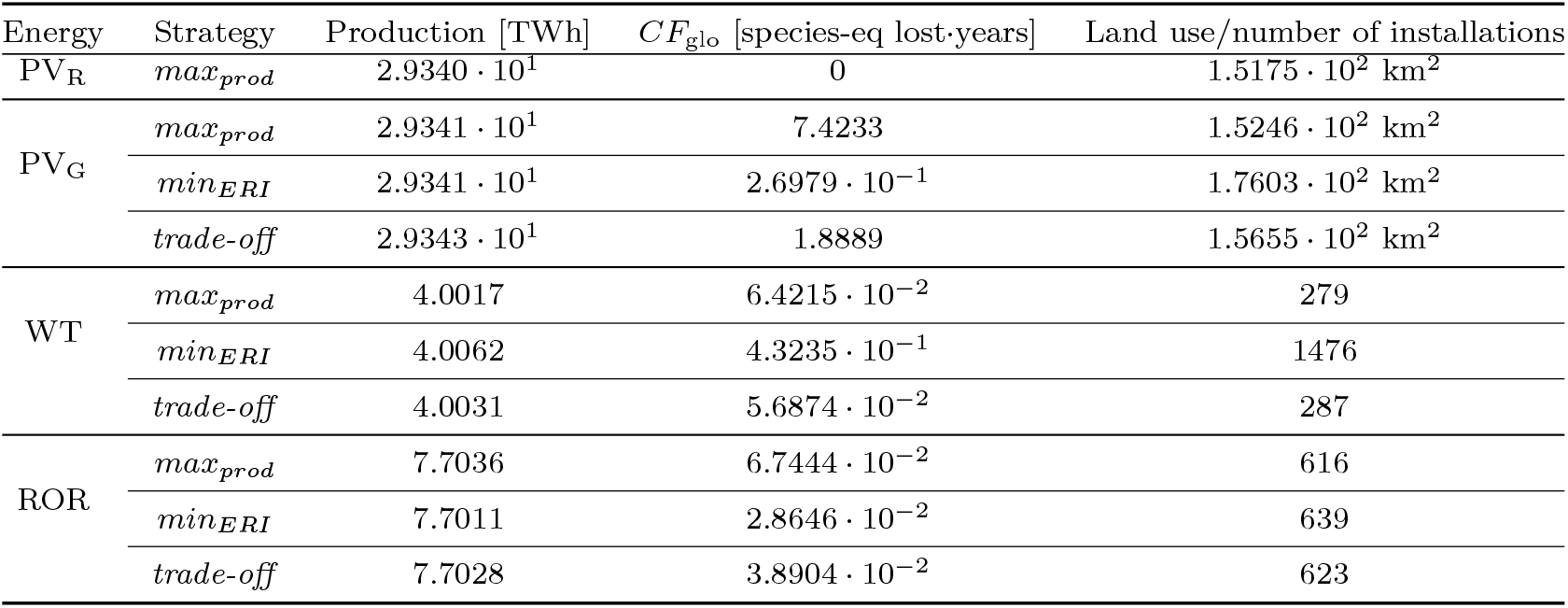
Summary of electricity production, biodiversity impact (*CF*_glo_), and infrastructure size (in area or number of units) for rooftop photovoltaic (PV_R_), ground-mounted photo-voltaic (PV_G_), wind turbines (WT), and small run-of-river hydropower (ROR) under the siting strategies *max*_*prod*_, *min*_*ERI*_, and *trade-off*.

### Maximizing production

First, we apply the *max*_*prod*_ siting strategy maximizing electricity production of each renewable technology. For rooftop PV, this strategy leads to the selection of approximately 236 thousand roofs (Figure B2a), covering a rooftop area of approximately 152 km^2^. The majority of these installations are situated in low-altitude regions (Figure B2b), with approximately 60 % concentrated between 400 m and 600 m in elevation and less than 2 % located above 1500 m. The installations are spread out across the country.

Ground-mounted PV installations leads to the selection of pixels covering a total land area of approximately 152 km^2^ (Figure B3a), and results in a national biodiversity impact of 7.42 species-eq lost·years. About 38 % of these installations are situated above 1500 m (Figure B3b). The panels are mostly located in agricultural and wooded areas, with about 59 % and 35 %, respectively, and the remaining 6 % cover unproductive land.

When siting WT to maximize the electricity production (Figure B3a), 279 WT are built resulting in a national biodiversity impact of approximately 6.42 × 10^−2^ species-eq lost·years (Figure B7a). The vast majority (90 %) is sited below 1500 m (Figure B7b). The installations are located in wooded (59 %) and agricultural (41 %) areas.

Favouring river sections with higher production potentials, 616 watercourse sections are selected to each host a ROR plant, resulting in a national biodiversity impact of approximately 6.74 × 10^−2^ species-eq lost·years. The hydropower sites are spread across the country Figure B11a, with half of the installations in low-altitude river reaches (49 % under 1500 m) Figure B11b.

### Minimizing extinction risks

The second siting strategy applied minimizes the extinction risk (Equation (*min*_*ERI*_)). This strategy is applied for the siting of all renewable technologies apart from rooftop PV, which we assume to have no biodiversity impact and thus an ERI of zero.

By prioritizing sites with the lowest ERI for the impacted species groups, the total area for ground-mounted PV increases by approximately 15 % to 176 km^2^ compared to the strategy maximizing production, but with a total biodiversity impact reduced by over 96 % to approximately 2.70 × 10^−1^ species-eq lost·years. The vast majority (96 %) of the sites are located below 1500 m (Figure B4a, Figure B4b). Further, a larger share of installations is sited on agricultural land (76 %), while wooded and unproductive areas are selected for only 23 % and 1 %, respectively.

Applied to WT, the siting strategy minimizing ERI fails to protect biodiversity and increases national impacts to 4.32 × 10^−1^ species-eq lost·years, that is almost 7 times higher than the solution siting of Equation (*max*_*prod*_). This is due to the selection of low productivity sites, thus increasing the total number of installed WT to meet the electricity generation target to 1476 WT, that is 5 times more installations compared to Equation (*max*_*prod*_). The WT are almost evenly located in both altitudes below (49 %) and above (51 %) 1500 m on agricultural, wooded, and unproductive land, accounting for 53 %, 38 %, and 9 %, respectively (Figure B8a, Figure B8b).

When applied to small ROR hydroelectric plants, the strategy minimizing ERI selects 639 sites, which corresponds to an increase by 4 % compared to Equation (*max*_*prod*_). However, the global impact is reduced of approximately 58 %, with 2.86 × 10^−2^ species-eq lost·years. Two thirds of the plants lie in high altitudes areas above 1500 m (Figures B12a and B12b).

### Trade-off strategy

Finally, we analyze siting decisions that balance electricity production and ERI by applying the *trade-off* strategy (Equation (*trade-off*)) to all renewable technologies except rooftop PV, as this technology is assumed to have no impact on biodiversity. The trade-off strategy uses a weighting factor *λ* to balance electricity production and ERI, generating Pareto-optimal siting configurations that meet the electricity production target. The selected trade-off solution is the non-dominated configuration closest to the ideal point (see Section 2.4).

The total land footprint of the renewable installations and the global CFs are shown for varying values of *λ* in Figure B5a for ground-mounted PV. As *λ* increases, the area requirement monotonically increases, whereas the biodiversity impact decreases sharply. An estimate of the Pareto front for the trade-off between land footprint and biodiversity impacts is estimated and represented in Figure B5b.

Using the least-distance-to-ideal method (see Section 2.4), we identify the siting configuration associated with the optimal trade-off weighting parameter 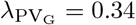 for ground-mounted PV. The spatial distribution of selected sites for the trade-off strategy is shown in Figure B6a and the associated altitude distribution is presented in Figure B6b. The tradeoff strategy results in a total land use of 157 km^2^, which is 3 % higher than *max*_*prod*_ but 11 % lower than *min*_*ERI*_ . The corresponding national biodiversity impact is reduced by 75 % compared to the *max*_*prod*_ strategy to 1.89 species-eq lost·years, but is 7 times higher than that of *min*_*ERI*_ . Low-altitude locations (below 1500 m) account for 65 % of the selected sites, while the remaining are situated above 1500 m elevation. The installations are predominantly located in agricultural areas, representing 77 % of the land selected, with the remaining share being located in wooded (19 %) and unproductive (4 %) areas.

The number of WT to deploy and the corresponding biodiversity impacts generated are evaluated for varying values of *λ* and shown in Figure B9a. As *λ* increases, the number of WT rises monotonically. For small *λ* values, the number of WT remains nearly constant, increasing only slightly from 279 at *λ* = 0 to 302 at *λ* = 0.6. Beyond this range, the number of WT increases sharply, reaching 1476 units at *λ* = 1. In parallel, for *λ* ≤ 0.8, the biodiversity impacts show limited variation, from 5.36 × 10^−2^ to 6.51 × 10^−2^ species-eq lost·years, followed by a pronounced rise at higher *λ* values, peaking at 4.33 × 10^−1^ species-eq lost·years for *λ* = 1. When priority is given to minimizing the extinction risk, turbine locations shift to less productive areas, requiring more WT to meet the production target. Lower extinction risks on a local level are overcompensated by higher infrastructure needs, ultimately resulting in increased ecological costs.

As with ground-mounted PV, we exclude dominated solutions to estimate the Pareto front (Figure B9b). In this case, we identify *λ*_WT_ = 0.48 as the trade-off strategy for WT, balancing infrastructure deployment and ecological impact. The corresponding selected pixels for wind energy installations are shown in Figure B10a, and their altitude distribution is presented in Figure B10b. In total, 287 WT are planned under this trade-off configuration, that is a slight increase of 3 % compared to *max*_*prod*_ and a substantial 81 % decrease compared to *min*_*ERI*_ . The trade-off configuration results in a global impact of 5.69 × 10^−2^ species-eq lost·years, which is lower than the impacts generated by both the *max*_*prod*_ and *min*_*ERI*_ siting strategies with 11 % and 87 % decreases, respectively. A total of 89 % of the WT are located below 1500 m. Wooded areas account for 63 % of the selected land, and the remaining 37 % are located in agricultural zones.

The number of small ROR plants and the associated biodiversity impacts are shown in Figure B13a for varying values of *λ*. The number of ROR hydroelectric plants increases with *λ*, while the global biodiversity impact decreases. The estimated Pareto front is shown in Figure B13b.

In this case, *λ*_ROR_ = 0.7 is the trade-off strategy for ROR plants siting. The strategy results in the selection of 623 river reaches for small hydropower installations, which is slightly higher (1 %) compared to the *max*_*prod*_ strategy, and slightly lower (−3 %) than the infrastructure requirement of the *min*_*ERI*_ strategy. The trade-off configuration induces a global impact of 3.89 × 10^−2^ species-eq lost·years, 42 % lower and 36 % higher compared to *max*_*prod*_ and *min*_*ERI*_, respectively. The spatial distribution of these river reaches is shown in Figure B14a, displaying the centroids of the reach polygons, and their altitude distribution is presented in Figure B14b. Installations located above 1500 m account for 62 % of the total.

The spatial distribution of the trade-off siting solutions of all renewable technologies, together with the relative shares of land cover types for solar and wind energy, are shown in Figure 1.

**Fig. 1.**
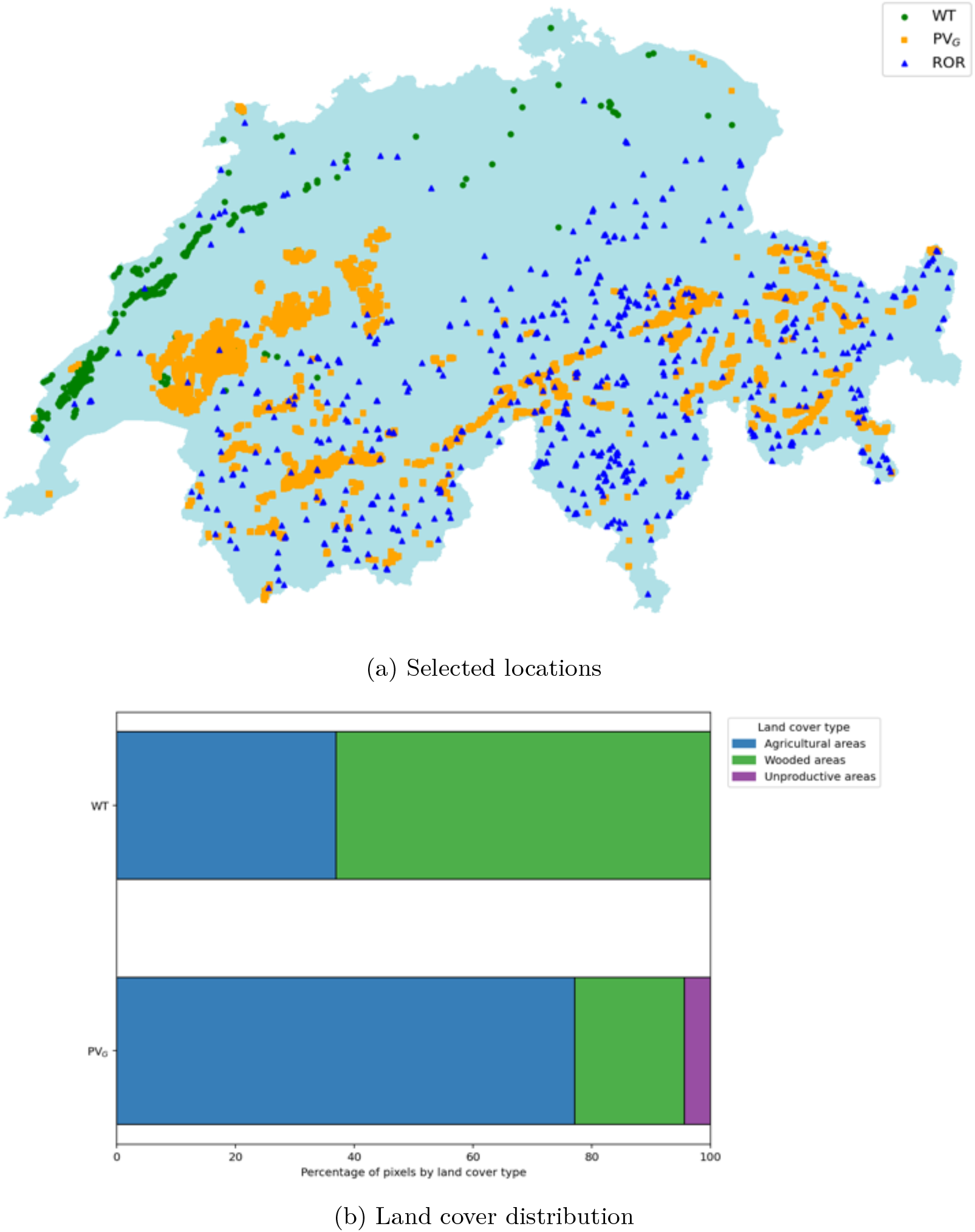
Locations of the trade-off siting solutions for ground-mounted photovoltaic (PV_G_), wind turbines (WT), and small run-of-river hydropower (ROR) (a), and their relative shares of land cover types for PV_G_ and WT (b). Unproductive areas correspond to unwooded, non-built areas that are unsuitable for cultivation.

### Biodiversity impact curves

In this section, the impacts on biodiversity of the different renewable energy technologies are assessed for increasing production targets (see Figure 2).

**Fig. 2.**
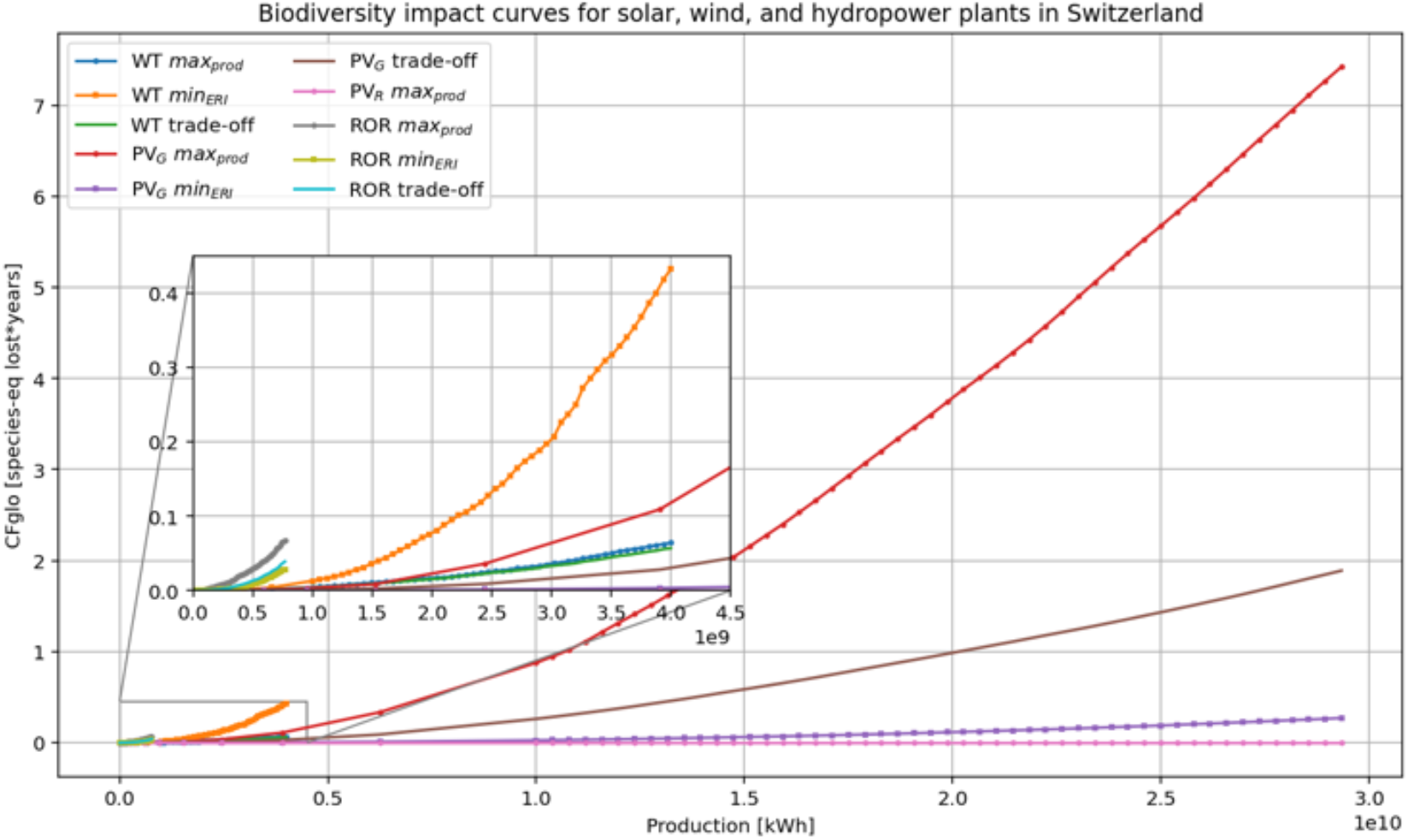
Biodiversity impacts of rooftop photovoltaic (PV_R_), ground-mounted photo-voltaic (PV_G_), wind turbines (WT), and small run-of-river hydropower (ROR) as a function of electricity production under the siting strategies *max*_*prod*_, *min*_*ERI*_, and *trade-off*.

Using the three siting strategies defined in Section 2.4, and selecting 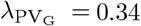 for the trade-off approach, the biodiversity impact curves corresponding to the strategies prioritizing areas of maximum energy potential (Equation (*max*_*prod*_)), areas with lower extinction risk for impacted species groups (Equation (*min*_*ERI*_)), and the trade-off between minimal land use and biodiversity impact are shown in Figure C15a for ground-mounted PV, and the associated areas used are depicted in Figure C15b.

Across all levels of solar energy conversion, the three siting strategies are ranked consistently in terms of biodiversity impact. The *max*_*prod*_ strategy yields the highest impacts, followed by the *trade-off* strategy, whereas the *min*_*ERI*_ strategy systematically results in the lowest impacts. Notably, the impacts associated with the *max*_*prod*_ strategy increase at a substantially steeper rate, highlighting its disproportionately stronger effect on biodiversity as production levels rise. When considering the area required for ground-mounted PV installations, the ranking of the three strategies remains consistent but becomes inverted: the *max*_*prod*_ strategy requires the smallest surface area, followed by the *trade-off* strategy, while the *min*_*ERI*_ strategy requires the largest area. The latter strategy also exhibits the steepest increase in required surface area as electricity production rises.

Similarly, we investigate the impacts and required infrastructure sizes of wind power for a production target varying between 0 up to 4 TW h (Figures C16a and C16b), which corresponds to the planned wind power production expansion between 2025 and 2050, as well as for small ROR hydropower plants up to the national expansion target of 0.77 TW h (Figures C17a and C17b). We use the optimal trade-off parameters *λ*_WT_ = 0.48 and *λ*_ROR_ = 0.7 for WT and ROR installations, respectively.

Figure 2 shows the cumulative biodiversity impacts measured as the global CFs, as a function of electricity production for all the technologies considered, under the different siting strategies. For reference, rooftop PV is included as a baseline scenario, and because this technology is assumed to have negligible land-use–related biodiversity effects [Katzner et al., 2013], its curve remains effectively at zero across all production levels. Comparing curves across the other energy types considered, ROR systems show the steepest impact gradients, with the highest biodiversity impacts over the entire production range considered. Their impacts increase rapidly at low electricity outputs and continues to rise steadily with increasing production. Ground-mounted PV plants globally yield the lowest biodiversity impacts compared to ROR and WT technologies, with an exception when considering the *max*_*prod*_ strategy. The latter used for ground-mounted PV, results in an impact that increases faster than WT curves for the *min*_*ERI*_ and *trade-off* approaches. Finally, WT globally exhibit the second-highest impact levels. Although their curves start with relatively similar values to ROR at low production, the rate of increase is less pronounced.

## 4 Discussion

In this study, we developed and applied a framework for siting infrastructure that incorporates biodiversity protection by using fine-scale datasets and accounting for a broad range of aquatic and terrestrial species groups. Rather than representing biodiversity as a fixed spatial constraint independent of infrastructure placement, the framework quantifies biodiversity impacts as a function of the selected siting configuration. Therefore, it captures how alternative siting strategies redistribute ecological pressure. In contrast, traditional spatial conservation planning tools [Ball and Possingham, 2000, Ball et al., 2009, Moilanen et al., 2005, Ciarleglio et al., 2009] generate relative conservation priorities but do not provide implementable infrastructure siting guidance [McIntosh et al., 2018]. Specifically, we assessed how deployment choices influence the ecological impacts of rooftop and ground-mounted PV, WT, and small ROR hydroelectric plants. Three siting strategies (Section 2.4) were evaluated and compared in terms of their biodiversity impacts and land footprint. For each renewable energy technology, the relevant impacted species groups were identified (Table 3), and detailed biodiversity maps were used to estimate site-specific extinction probabilities. The latter were then combined with CFs to quantify national level biodiversity impacts of renewable energy infrastructure (Section 2.1). This integration of highresolution data extends LCA-based biodiversity metrics, which are typically applied at coarse spatial scales or global averages [De Baan et al., 2013, Verones et al., 2013, May et al., 2020, Chaudhary et al., 2015, Scherer et al., 2023], toward a spatially explicit decision-support context suitable for infrastructure planning. Furthermore, to our knowledge, the inclusion of such a broad range of taxonomic groups represents a novel contribution, as previous LCA-based studies generally focus on a limited set of species groups [Damiani et al., 2023]. Although the empirical results presented here are specific to Switzerland, the methodological framework is transferable to other regions provided that sufficiently resolved data are available.

The scalability of rooftop PV is constrained by the limited availability of suitable rooftop area and its lower production potential compared to ground-mounted PV. However, given its negligible biodiversity impact, it serves as a reference baseline in our study. For a given energy type, we observe a consistent ordering of the three siting strategies for increasing electricity production targets, both in terms of infrastructure requirements and biodiversity impacts (Figure 2). The persistence of these patterns highlights that biodiversity-aware siting leads to systematic and predictable changes in infrastructure distribution and biodiversity impacts, supporting transparent evaluation of planning trade-offs that would not be captured by coarse-scale approaches or limited-taxon studies [Zelm et al., 2011, Geyer et al., 2010, de Baan et al., 2015, Verones et al., 2013]. Importantly, this stable ranking of strategies for a given technology likely reflects structural trade-offs between energy potential and ecological sensitivity, and may therefore extend beyond the Swiss context.

The biodiversity impacts of ground-mounted PV and ROR power plants are highly sensitive to siting decisions. Notably, reductions in biodiversity impact are achieved at the cost of increasing area demand. The max_*prod*_ strategy, which prioritizes pixels with the highest energy potential, yields the smallest installation footprint (Figures C15b and C17b) but the highest biodiversity impacts for ground-mounted PV and ROR power plants (Figures C15a and C17a). In contrast, strategies that incorporate extinction risk information of impacted species groups can yield significant reductions in biodiversity impact, with only modest increases in land use or infrastructure requirements. Explicitly accounting for species-group vulnerabilities reshapes optimal deployment patterns, revealing trade-offs that are not apparent when energy performance alone guides siting [May et al., 2021, Dorber et al., 2020]. This observation underscores the importance of integrating biodiversity considerations into renewable energy system planning. However, for wind power, prioritizing areas of low extinction risk results in the selection of areas with low availability, dramatically increasing the number of turbines required to meet production targets (Figure C16b). Consequently, prioritizing low local extinction risk paradoxically increases the overall impact (Figure C16a) due to substantially higher infrastructure requirements. This demonstrates that siting strategies based solely on local biodiversity considerations can backfire at the system scale. Instead, effective siting strategies must balance availability and biodiversity impacts on a system level, with trade-off configurations offering lower impacts for only marginally higher infrastructure needs.

Across all technologies, clear spatial patterns emerge when comparing the three siting strategies on our case study. The max_*prod*_ approach systematically concentrates installations in areas with high energy potential but also higher environmental sensitivity: low-elevation forested zones for wind, and substantial shares of wooded land for ground-mounted PV. In contrast, the min_*ERI*_ strategy shifts deployments toward less sensitive landscapes, typically agricultural areas and, for wind and ROR hydro, higher elevations. Trade-off solutions generally fall between these extremes but tend to resemble one or the other depending on the technology: for wind, the compromise remains close to max_*prod*_, whereas for ground-mounted PV it aligns more strongly with the ERI-oriented pattern, strongly favouring agricultural land. These spatial contrasts illustrate how biodiversity-informed siting can redirect infrastructure toward landscapes where ecological pressure is lower without eliminating energy potential, a capability that prior studies could not offer [May et al., 2021, Dorber et al., 2020].

In real-world siting decisions, additional criteria beyond plant availability and biodiversity impact must be considered, including technical constraints related to accessibility and grid connection as well as issues of social acceptance [Huber et al., 2017]. In the maximum production and trade-off strategies, installations tend to be heavily clustered in a limited number of regions, where a higher production potential of renewable plants is expected. Specifically, southern regions with higher solar irradiation account for a substantial share of the ground-mounted PV installations, while most additional small ROR hydroelectric plants are concentrated in mountainous regions with suitable hydrological conditions. Such spatial clustering of infrastructure can exacerbate social acceptance challenges, which remain a critical non-technical barrier to the deployment of renewable energy [Bogdanov et al., 2019] and therefore must be included in siting decisions. The modular structure of the framework allows additional spatial and socio-technical criteria to be incorporated, supporting more comprehensive planning analyses [Moilanen et al., 2009, Pressey et al., 2007, Kienast et al., 2017].

When interpreting the results of this study, several limitations should be considered. First, impacts on biodiversity are highly region-specific and require detailed ecological surveys that are hard to generalize. Here, we calculate the ERI using high-resolution species distribution data for Switzerland. This indicator does not account for species migration within or across borders. However, species that disappear in Switzerland may persist elsewhere and potentially recolonize from neighbouring regions. Consequently, our extinction metrics should not be interpreted as measures of irreversible global loss.

Additionally, our analysis assumes static species distributions, neglecting ecological dynamics such as dispersal or climate-driven range shifts. This introduces uncertainty in the estimated biodiversity impacts, depending on species mobility and habitat connectivity.

Although species richness is a widely used metric for measuring biodiversity, it captures only one facet of biological diversity. Other dimensions, such as genetic, functional, or ecosystem diversity, are not represented in our indicators, which may oversimplify biodiversity responses to energy development. While this study focuses on species richness, future multi-criteria siting approaches should incorporate these additional components.

Finally, our assessment focuses on local land-use–related impacts occurring during the operational phase of renewable energy infrastructure. Upstream and downstream processes of the life cycle, such as construction, manufacturing, and waste treatment are intentionally excluded as may occur in other geographies with different ecological sensitivities. These life cycle stages can also pose risks to biodiversity in those locations.

These limitations suggest that our results represent a partial yet informative estimate of the overall biodiversity impacts associated with renewable energy expansion. While our dataset provides a detailed and spatially explicit assessment within Switzerland, comparable data are often unavailable at similar resolution in other countries, limiting the direct transferability of our approach. Therefore, the numerical impact levels and optimal parameter values reported here should be interpreted as context-specific rather than globally applicable. Nevertheless, the framework presented here provides a foundation for evaluating trade-offs between energy infrastructure deployment and local biodiversity conservation, and offers a basis for future refinement as more detailed ecological and life-cycle data become available, especially at broader geographical scales.

## 5 Conclusion

One essential strategy to reach net-zero emissions is to shift away from fossil fuels in favour of a substantial expansion of renewable energy capacity. Yet, this transition must be designed to avoid adding pressure on ecosystems and accelerating biodiversity loss, which would further degrade the benefits humans get from nature. This study presents a transferable framework for assessing and mitigating biodiversity impacts associated with infrastructure deployment. Although tailored here to renewable energy technologies, namely rooftop and ground-mounted PV, WT, and ROR hydroelectric plants, and demonstrated using Switzerland as a case study, the approach is designed to be applicable to other infrastructure types and geographical contexts.

Building on metrics initially designed for LCA integration, the framework integrates high-resolution species distribution data, installation-specific impacted species groups, and region-specific species vulnerabilities, considering 20 impacted species groups based on an extensive literature review. Combining this with spatially explicit production potential data of the considered infrastructure, we quantify biodiversity impacts under alternative siting strategies for projected renewable energy deployment in 2050.

Our results show that siting choices greatly influence biodiversity outcomes. A strategy that maximizes electricity production concentrates infrastructure in areas of high potential but also potentially high ecological sensitivity, leading to disproportionately high biodiversity impacts, particularly for ground-mounted PV and ROR plants. Conversely, strategies that minimize local extinction risk may shift deployment toward areas with lower generation potential, increasing installation areas. For wind energy, such strategies paradoxically substantially increase overall biodiversity impact. Trade-off strategies consistently offer more balanced outcomes, reducing ecological impacts relative to pure production maximization while avoiding the excessive infrastructure requirements of strict extinction risk minimization.

The effectiveness of these strategies is technology-dependent, demonstrating that a single cross-technology siting approach is not sufficient. Instead, the decision criteria should be technology-specific and explicitly account for biodiversity considerations. More importantly, our findings show that integrating biodiversity information into early-stage planning enables energy systems to be designed in ways that can significantly reduce harm to ecosystems.

Overall, the proposed framework provides a robust foundation for sustainable infrastructure planning and can be further refined or adapted to other regions. Moreover, the biodiversity impact functions can be directly integrated into energy system models, enabling dynamic and scenario based biodiversity-friendly planning. Future research could further reduce ecological impacts by exploring the large-scale potential of hybrid systems, integrating different renewable energy sources, such as leveraging existing hydropower infrastructure to exploit PV potential.

## 6 Associated content

### Data and code availability

The data and code generated in this study will be publicly available upon publication.

### Supporting Information

The Supporting Information will be available upon publication.

## 7 Acknowledgements

The authors thank Jérôme Dujardin for providing valuable maps of capacity factors for different types of wind turbines in Switzerland. They also thank Stefanie Hellweg, Stephan Pfister, and their team for their constructive feedback at an earlier stage of this work. The authors gratefully acknowledge funding from the Joint ETH-Initiative SPEED2ZERO. The project conducts research, develops tools, creates action plans, and implements technologies to support a sustainable transformation in Switzerland. A transformation that meets international and national climate targets ensures a resilient energy supply, allowing biodiversity to regain its richness. SPEED2ZERO received support from the ETH-Board under the Joint Initiatives scheme. S.M. and G.W. acknowledge support from the Swiss National Science Foundation (SNSF) under grant no. PZ00P2 202117.

## 8 Author information

### Author contributions

Marie-Ange Dahito: Conceptualization, Methodology, Software, Formal analysis, Investigation, Data curation, Visualization, Writing - original draft, Writing - reviewing & editing.

David Yang Shu: Conceptualization, Writing - reviewing & editing. Gabriel Wiest: Conceptualization, Writing - reviewing & editing.

Stefano Moret: Conceptualization, Writing - reviewing & editing. Tobias Wechsler: Data curation, Writing - reviewing & editing.

Loïc Pellissier: Conceptualization, Funding acquisition, Supervision, Writing - reviewing & editing.

## Appendix A

Production potentials and extinction risks

**Fig. A1.**
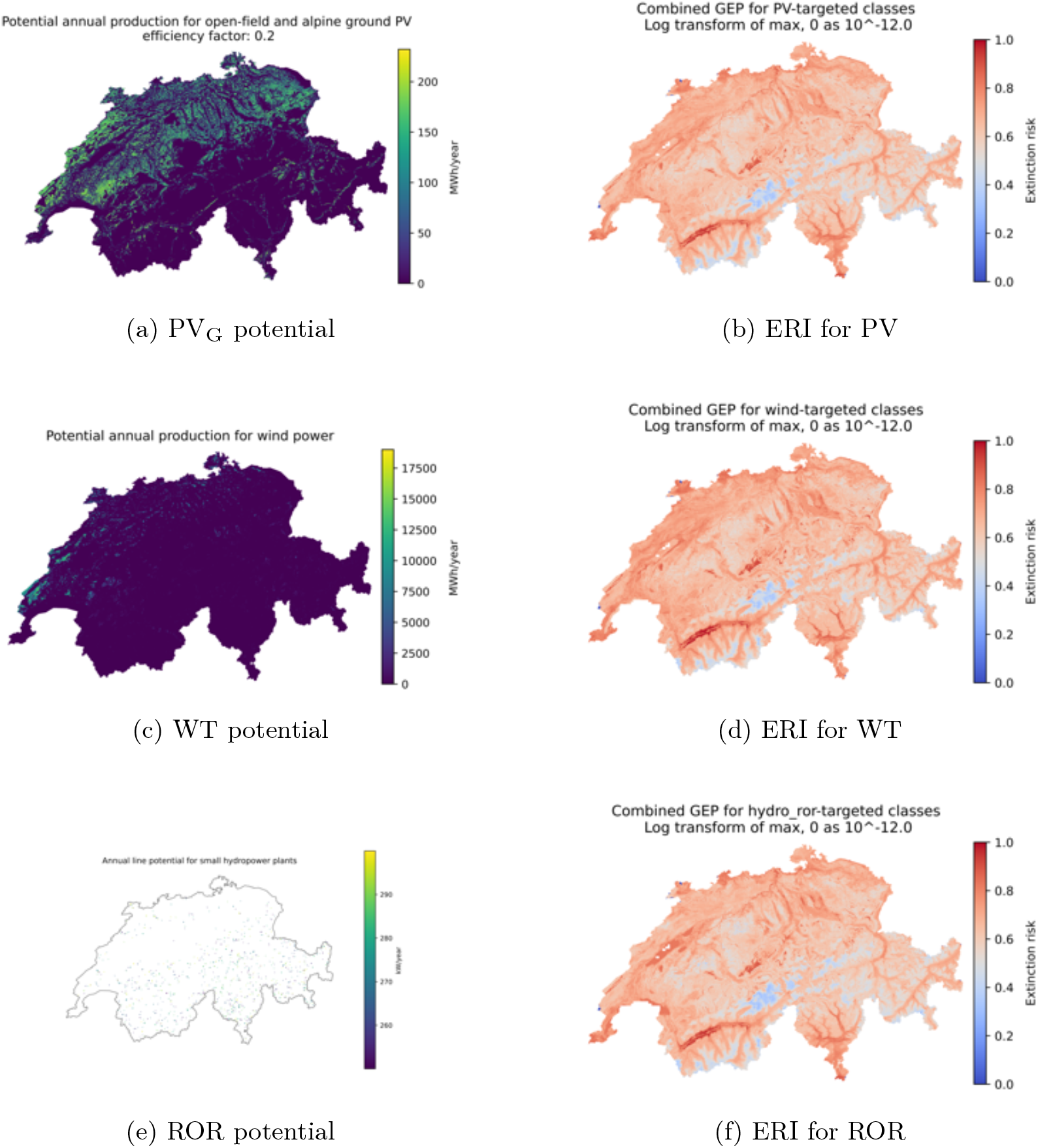
Maps of production potential for ground-mounted photovoltaic (PV_G_), wind turbines (WT), and small run-of-river hydropower (ROR) alongside the extinction risk indicator (ERI) values of species affected by the respective energy systems.

## Appendix B

Selected renewable energy sites and the associated national biodiversity impacts

### B.1 Rooftop photovoltaic

**Fig. B2.**
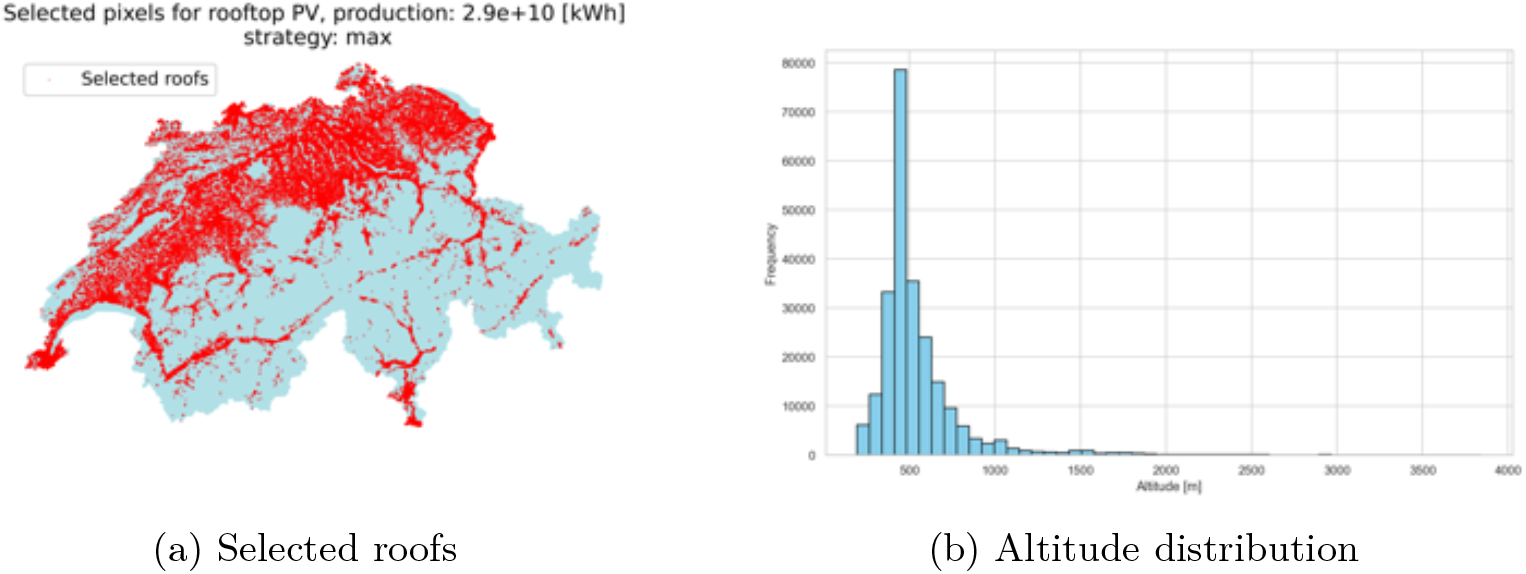
Selected roofs and their altitude distribution for rooftop photovoltaic production using strategy *max*_*prod*_.

### B.2 Ground-mounted photovoltaic

**Fig. B3.**
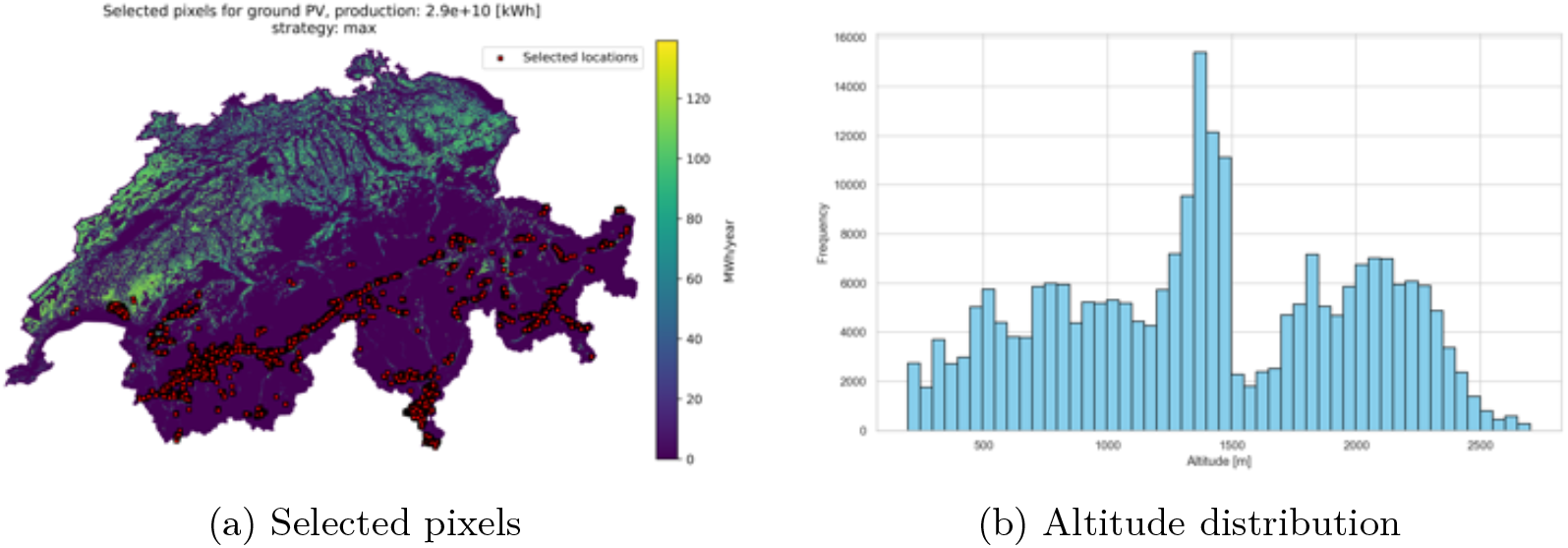
Selected pixels and their altitude distribution for ground-mounted photovoltaic production using strategy *max*_*prod*_.

**Fig. B4.**
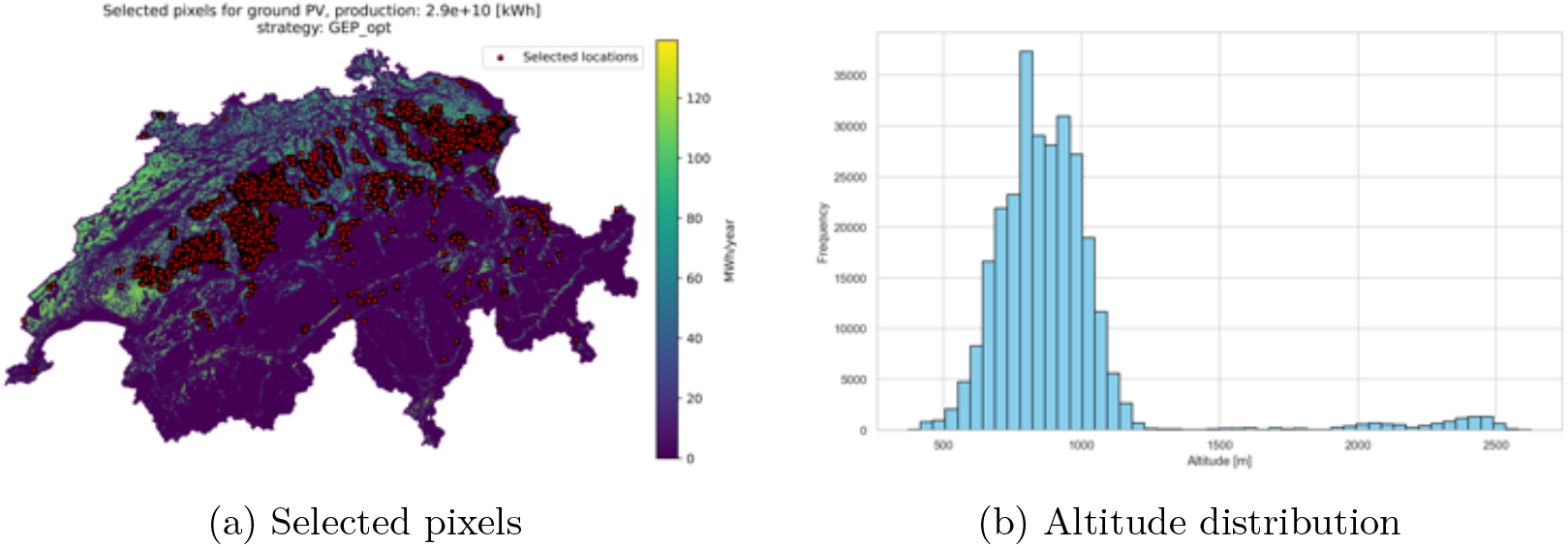
Selected pixels and their altitude distribution for ground-mounted photovoltaic production using strategy *min*_*ERI*_.

**Fig. B5.**
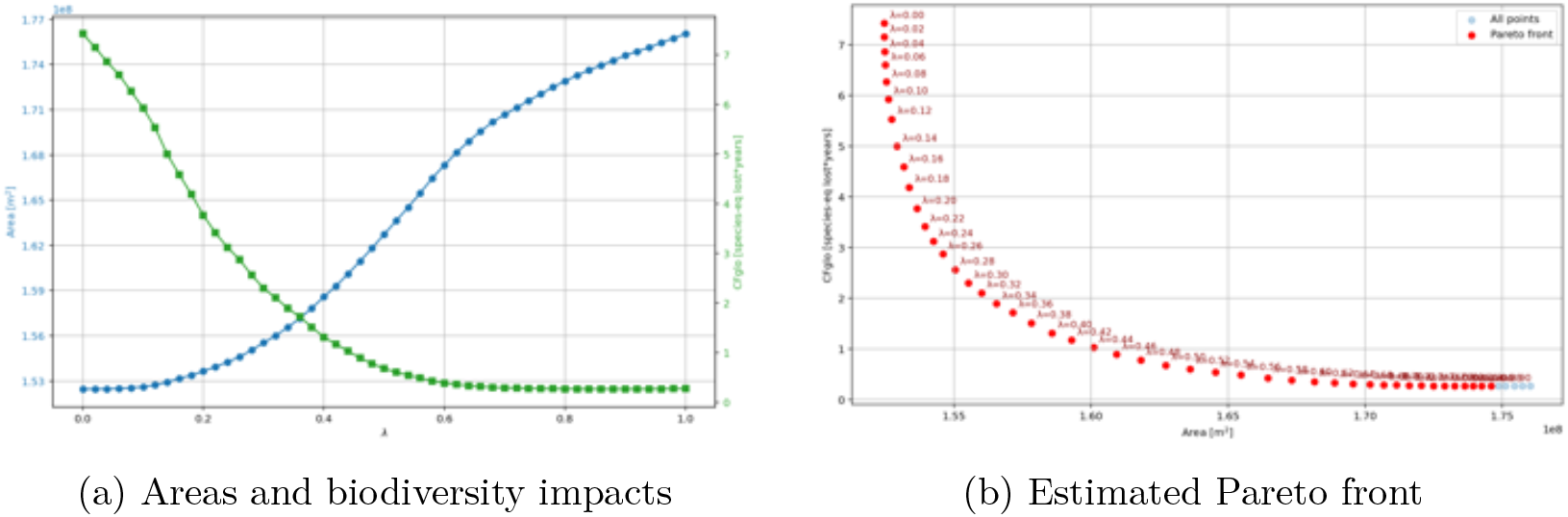
Areas required for ground-mounted photovoltaic panels and associated biodiversity impacts (a), and estimated Pareto front (b) across varying *λ* values under the strategy *trade-off*.

**Fig. B6.**
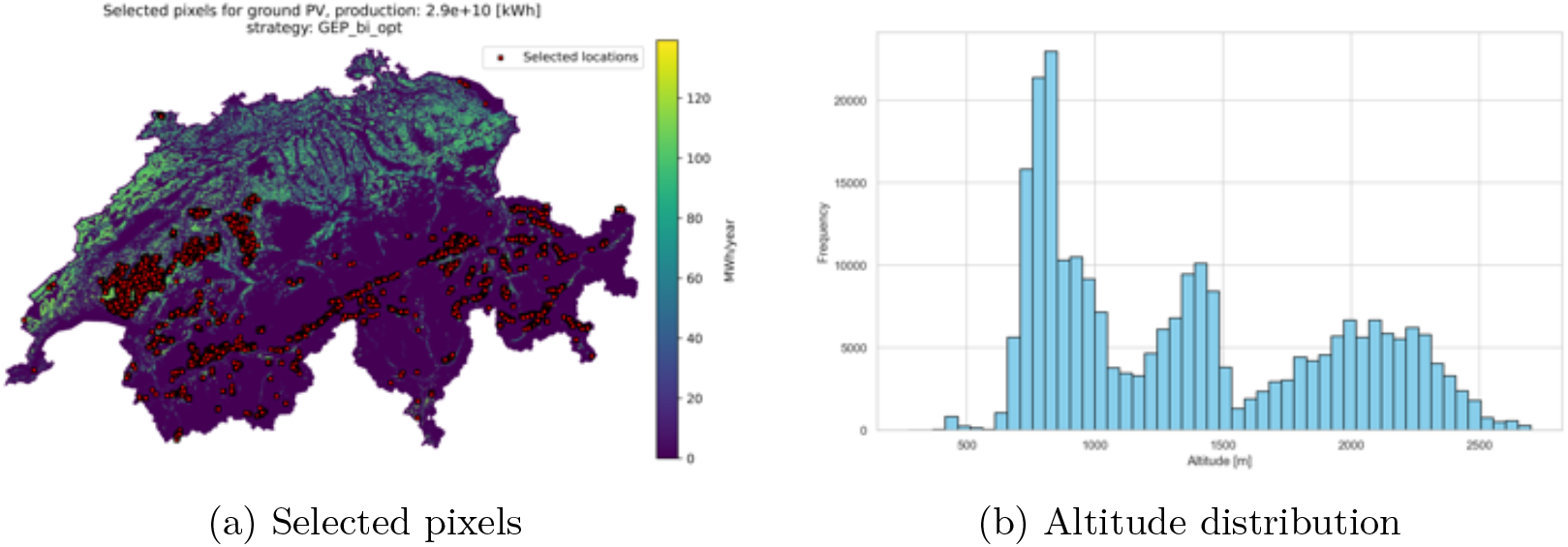
Selected pixels and their altitude distribution for ground-mounted photovoltaic production using strategy *trade-off*.

### B.3 Wind power

**Fig. B7.**
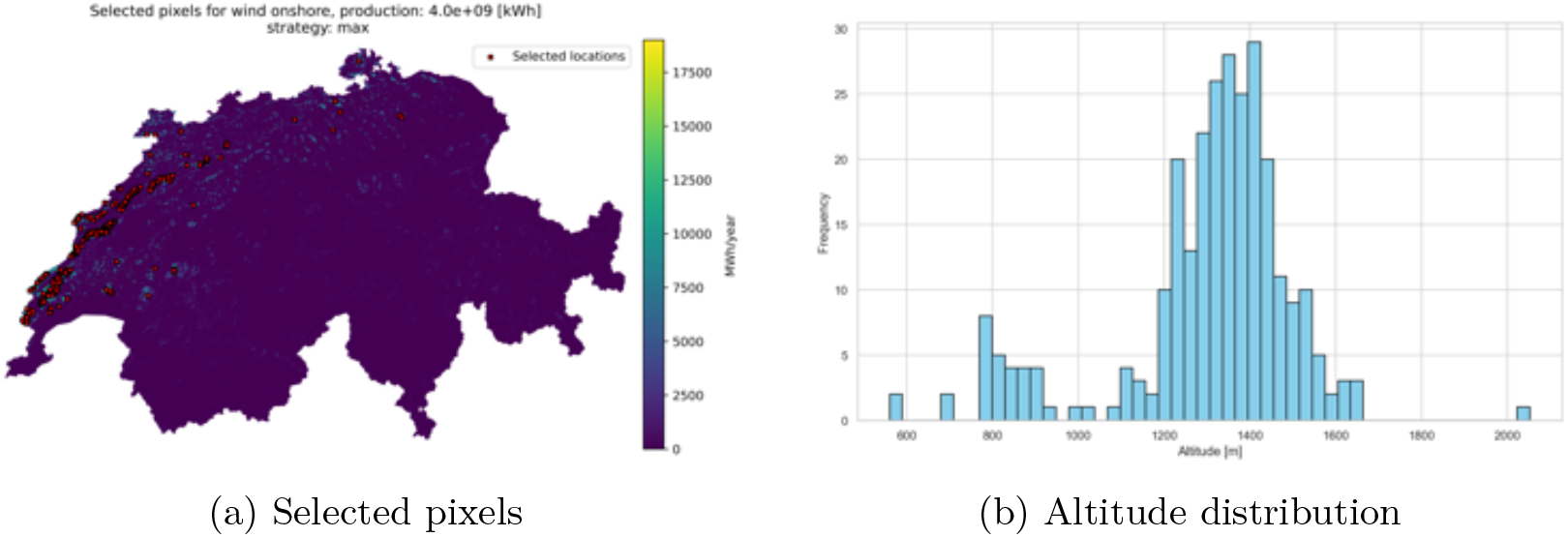
Selected pixels and their altitude distribution for wind power production using strategy *max*_*prod*_.

**Fig. B8.**
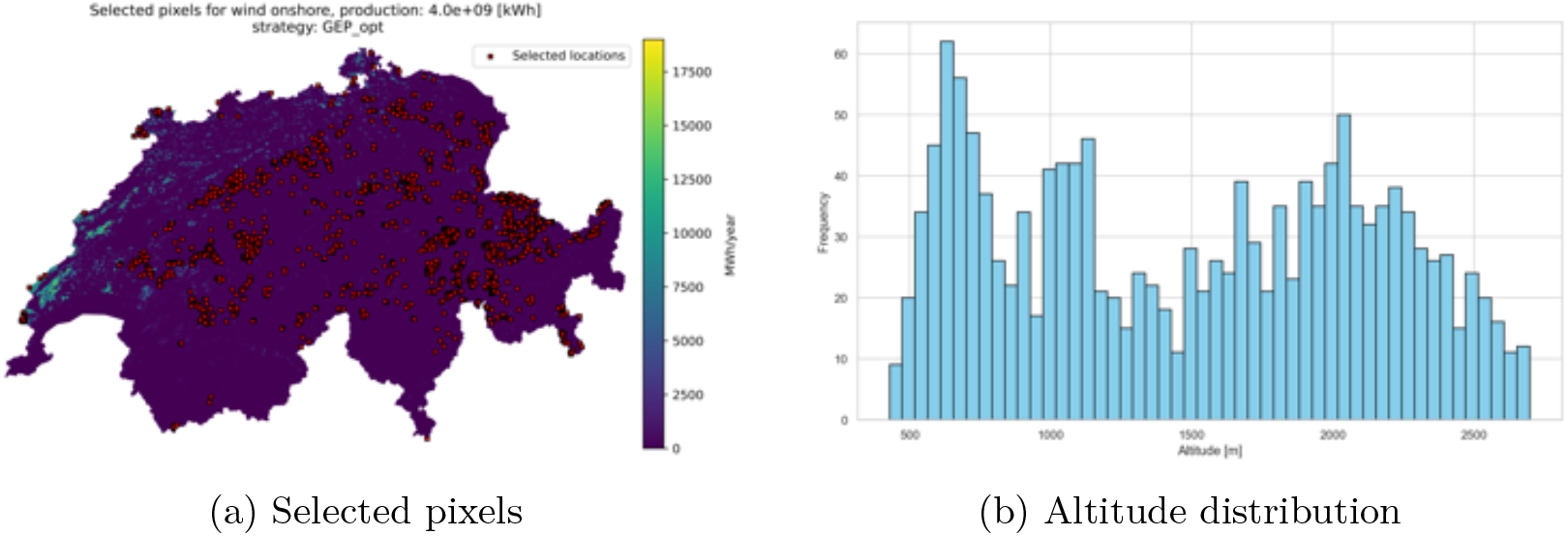
Selected pixels and their altitude distribution for wind power production using strategy *min*_*ERI*_.

**Fig. B9.**
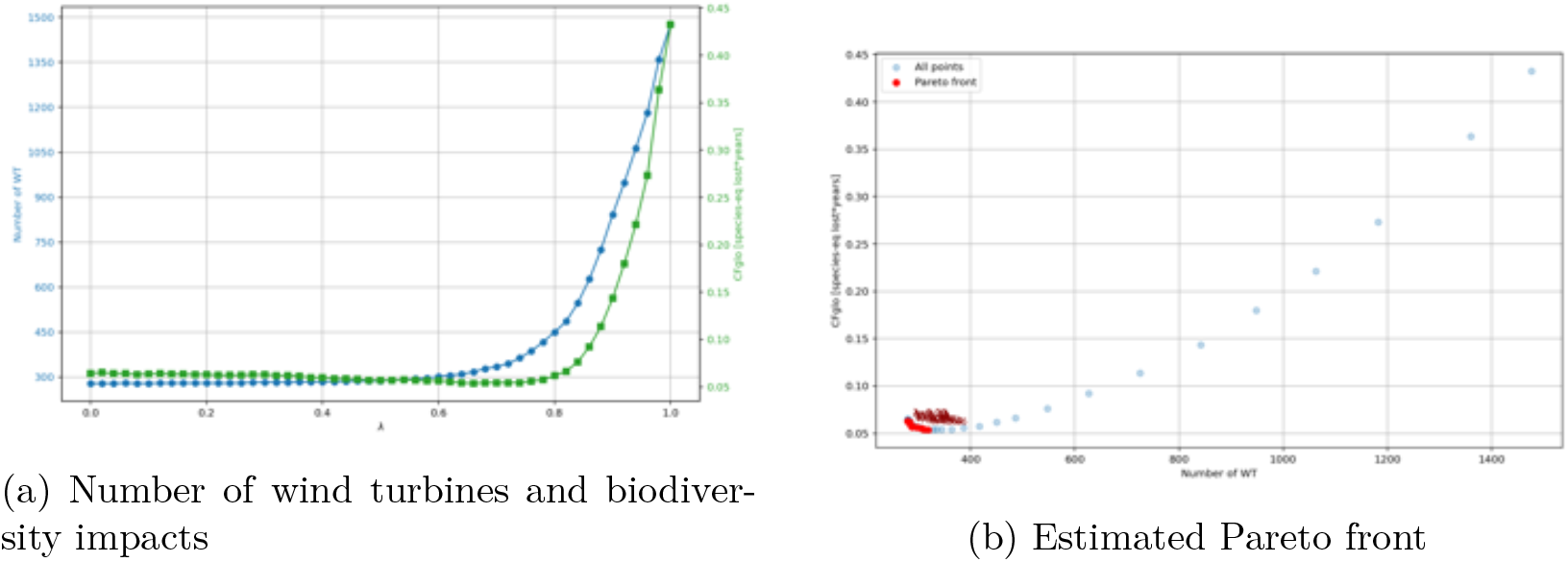
Number of wind turbines and associated biodiversity impacts (a), and estimated Pareto front (b) across varying *λ* values under the strategy *trade-off*.

**Fig. B10.**
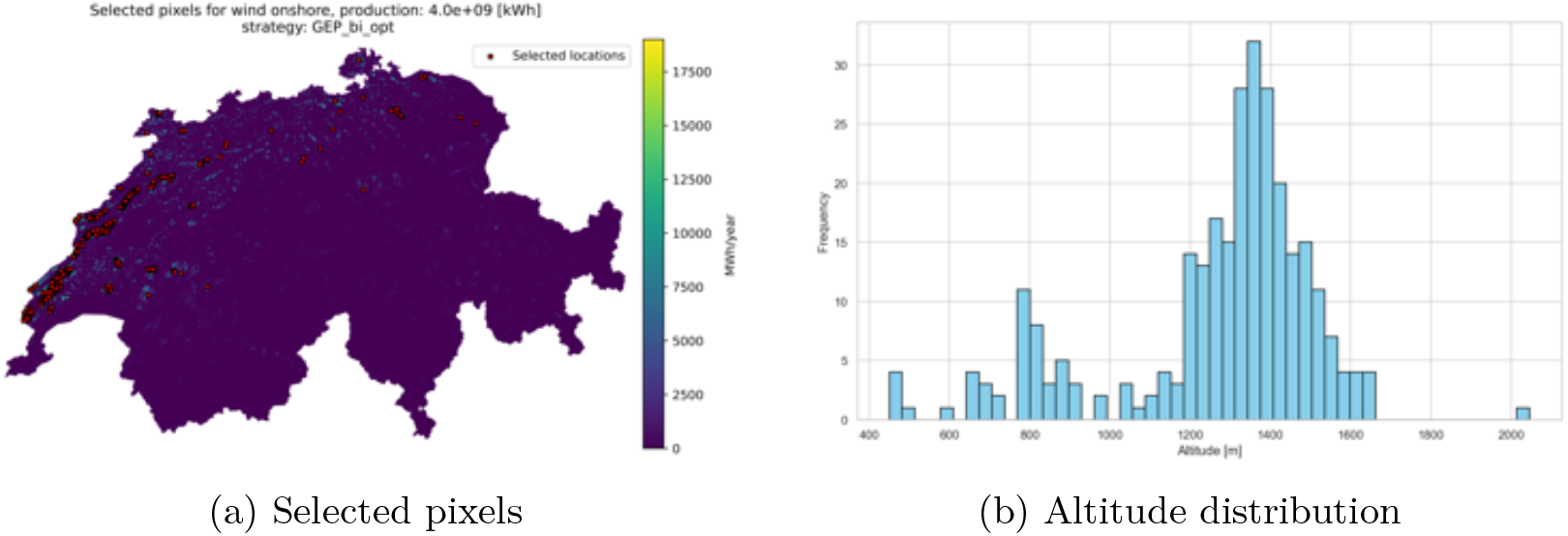
Selected pixels and their altitude distribution for wind power production using strategy *trade-off*.

### B.4 Small hydropower

**Fig. B11.**
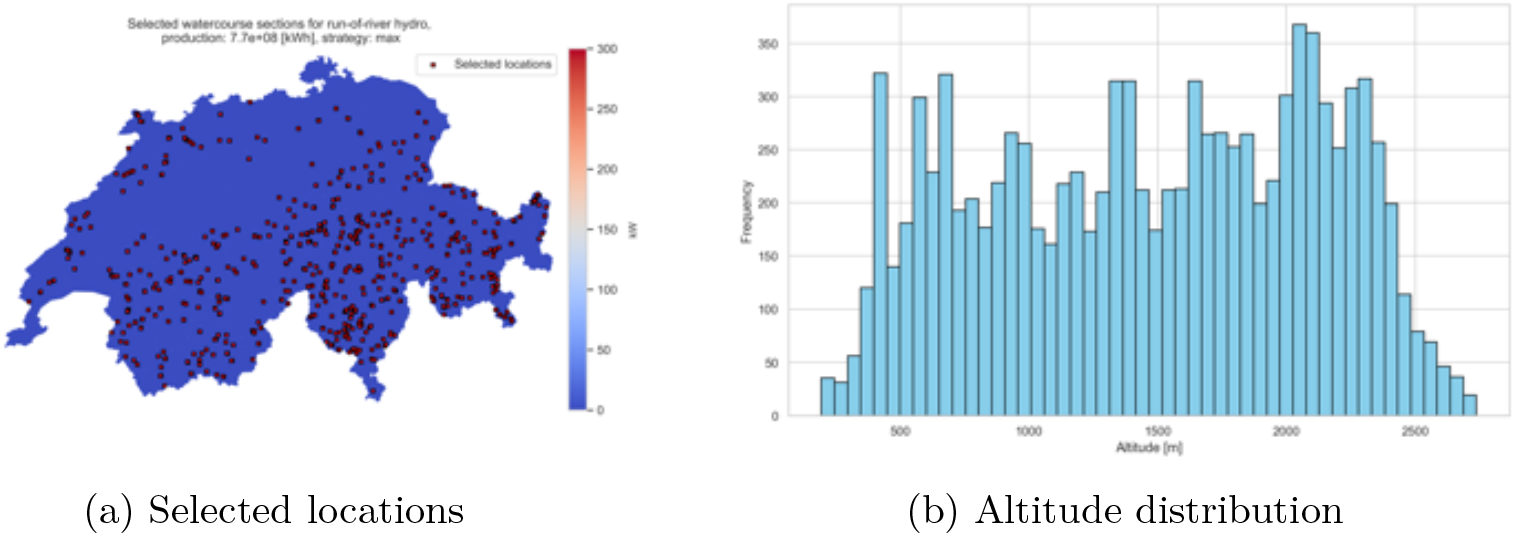
Selected locations (computed as the centroids of the river section polygon geometries) (a) and their altitude distribution (b) for small run-of-river hydropower production using strategy *max*_*prod*_.

**Fig. B12.**
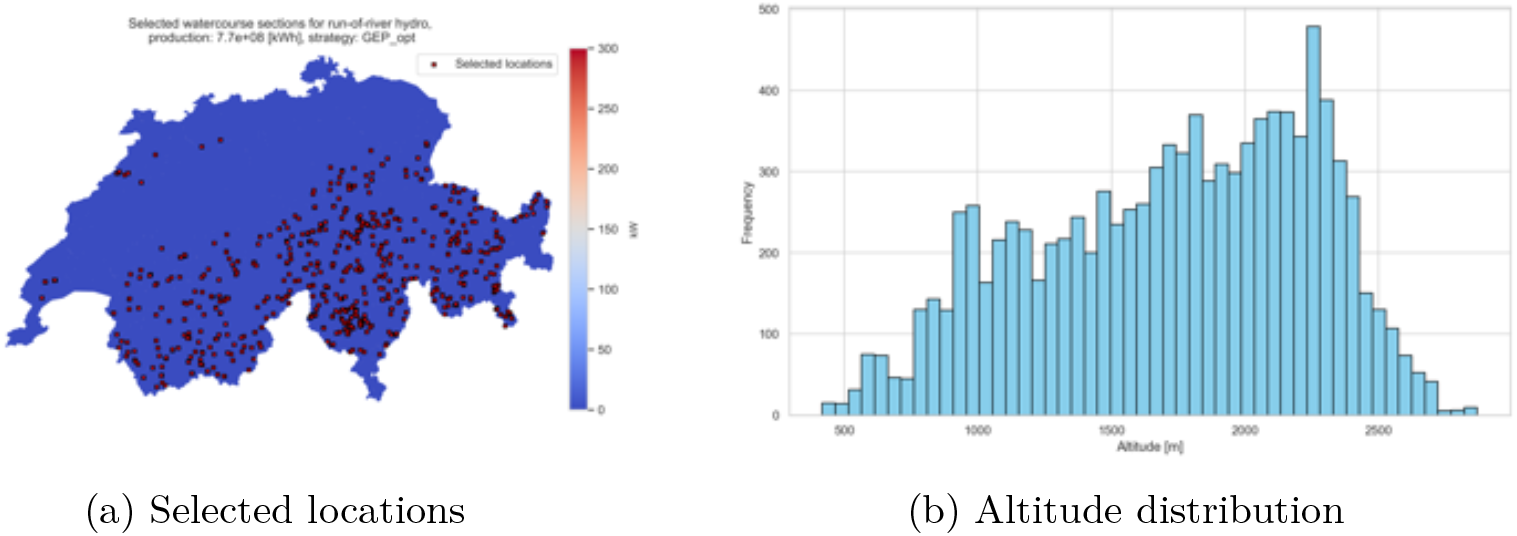
Selected locations (computed as the centroids of the river section polygon geometries) (a) and their altitude distribution (b) for small run-of-river hydropower production using strategy *min*_*ERI*_.

**Fig. B13.**
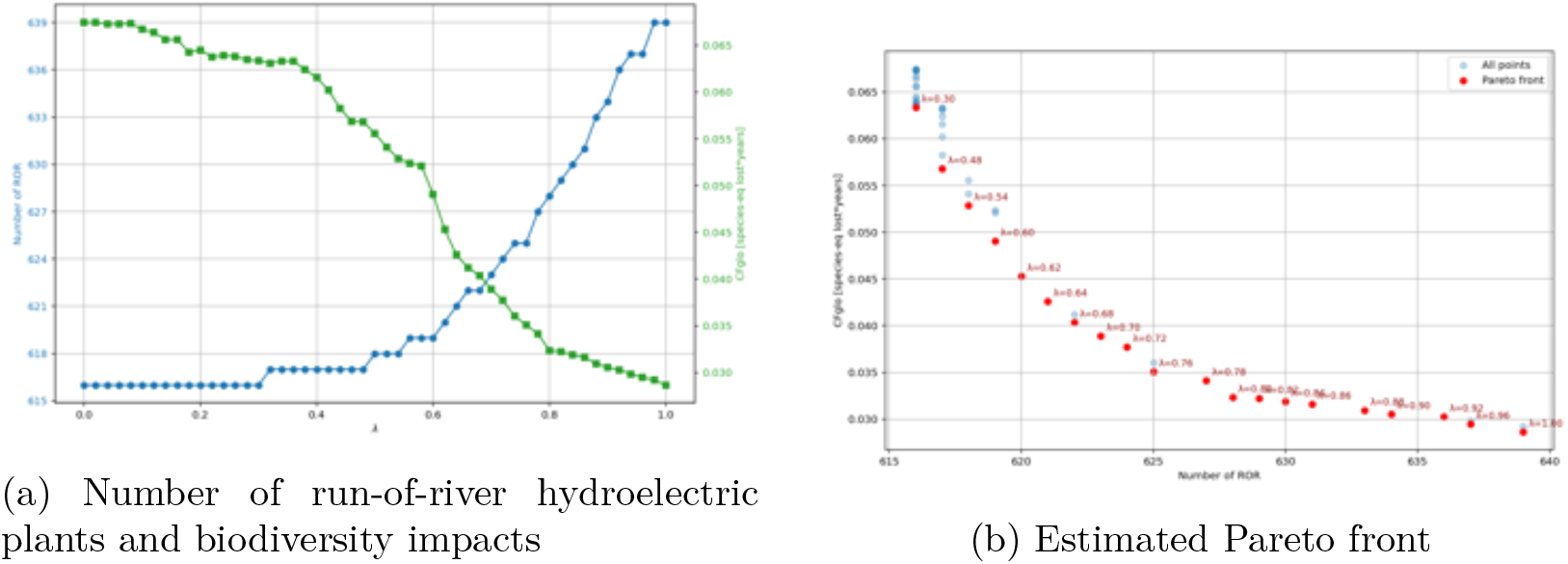
Number of small run-of-river plants and associated biodiversity impacts (a), and estimated Pareto front (b) across varying *λ* values under the strategy *trade-off*.

**Fig. B14.**
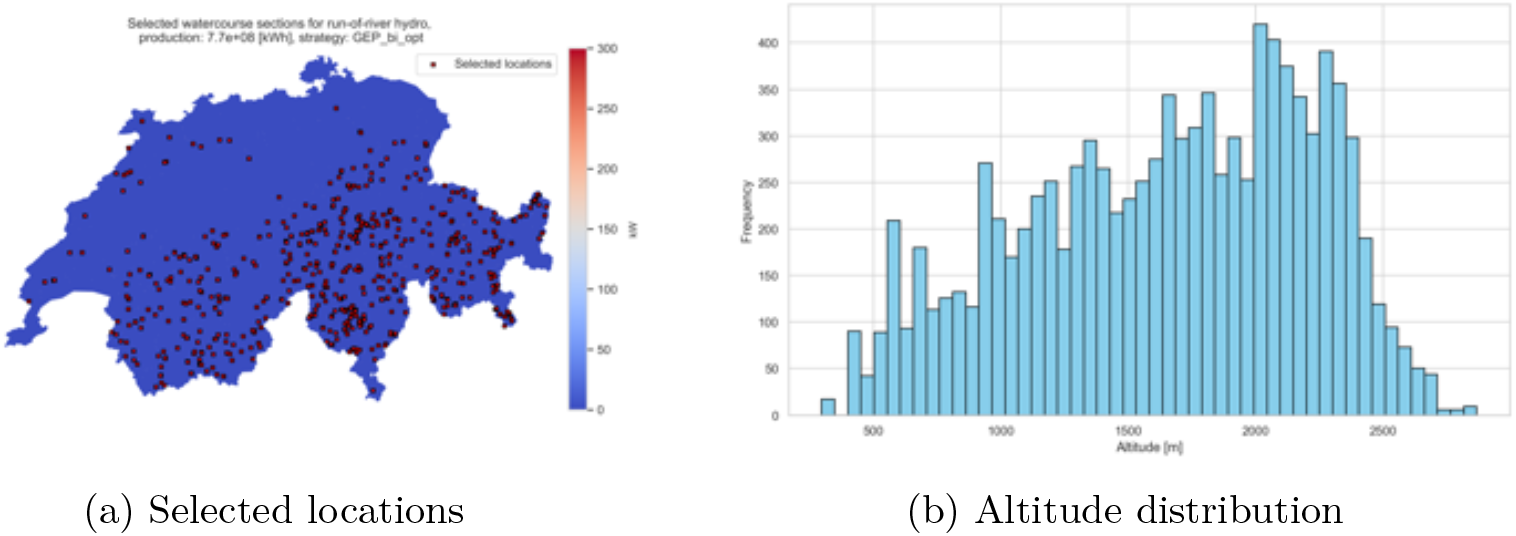
Selected locations (computed as the centroids of the river section polygon geometries) (a) and their altitude distribution (b) for small run-of-river hydropower production using strategy *trade-off*.

## Appendix C

Biodiversity impact curves

### C.1 Ground-mounted photovoltaic

**Fig. C15.**
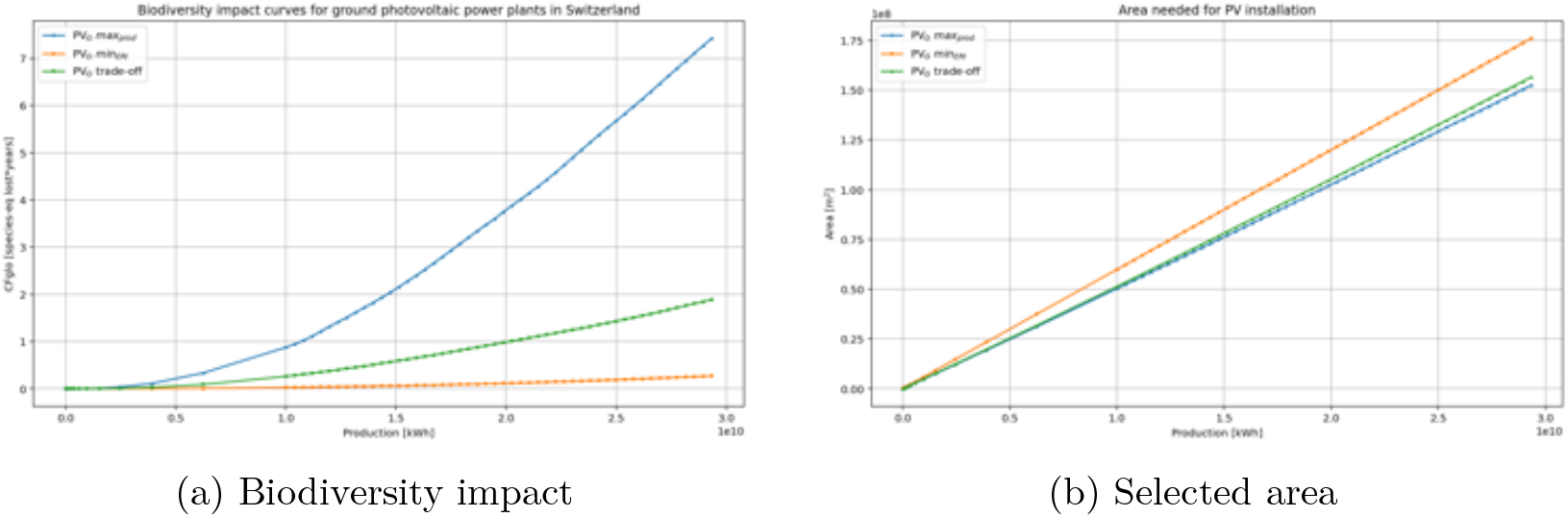
Biodiversity impact (a) and total selected area (b) for ground-mounted photovoltaic (PV_G_) installations, as a function of the production target, under strategies *max*_*prod*_, *min*_*ERI*_, and *trade-off*.

### C.2 Wind power

**Fig. C16.**
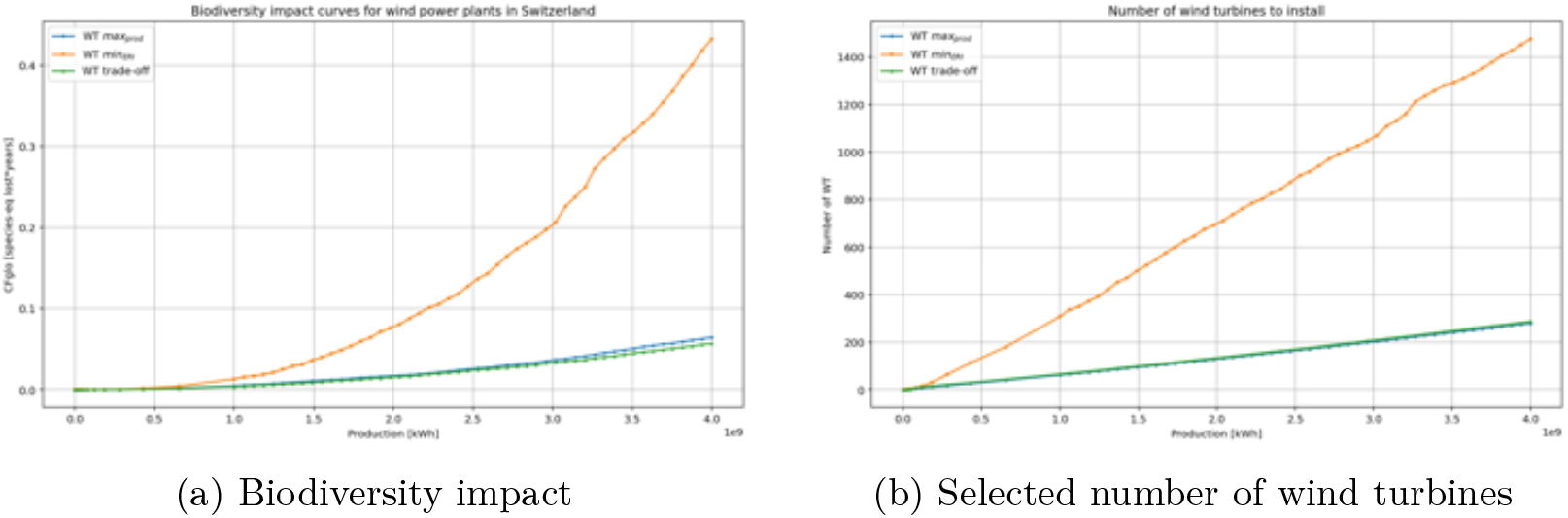
Biodiversity impact (a) and total number of installations (b) for wind turbines (WT), as a function of the production target, under strategies *max*_*prod*_, *min*_*ERI*_, and *trade-off*.

### C.3 Small hydropower

**Fig. C17.**
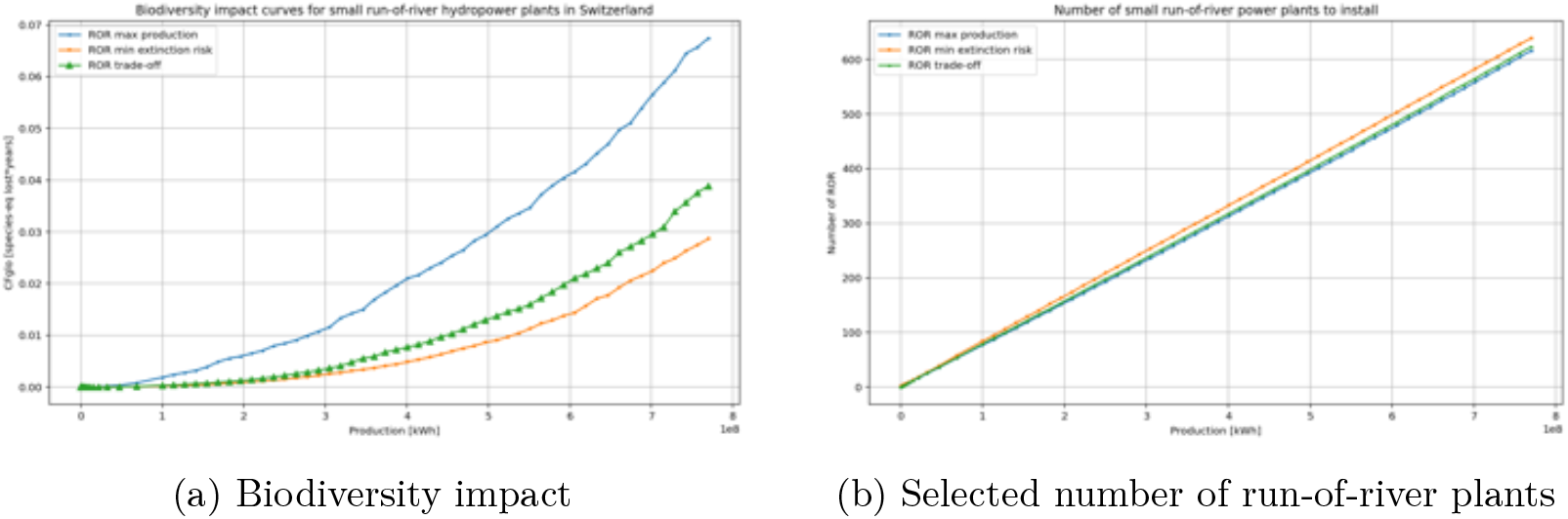
Biodiversity impact (a) and total number of installations (b) for small run-of-river hydropower (ROR), as a function of the production target, under strategies *max*_*prod*_, *min*_*ERI*_, and *trade-off*.

